# Conformational Switching between ON and OFF States Proceeds through Multiple Pathways in the SAM-III Translational Riboswitch

**DOI:** 10.64898/2026.09.10.749994

**Authors:** Dibyendu Mondal, Sabyasachi Paul Chowdhury, Govardhan Reddy

## Abstract

Gene regulation by translational riboswitches relies on repeated and reversible conformational switching between ON (*apo*) and OFF (*holo*) states. The switching mechanism often involves significant structural rearrangement, and the role of the metal ions in this mechanism is unclear. Using molecular dynamics simulations, we studied the switching transition of a translational riboswitch that regulates the concentration of Sadenosyl methionine (SAM), the ‘universal methyl donor’. We show that Mg^2+^ ions are essential for stabilizing the ON-state, and the three-way junction exhibits a ‘breathing-like’ dynamic interconversion between open and closed modes. The riboswitch has a low population of a *holo*-like READY (R) state with a partially organized SAM-binding pocket prior to ligand binding. Multiple intermediates are observed in the switching landscape, contributing to the gradual stepwise structural transition between the ON and R states, and proceeding through multiple competing pathways. Interestingly, Mg^2+^ ions influence the dominant pathway for the ON to R state transition, whereas the reverse transition shows a weak dependence. These results provide broad insights into the function of translational riboswitches, which repeatedly and rapidly respond to changes in metabolite concentrations, in contrast to transcriptional riboswitches.

## Introduction

Bacterial genetic regulation relies on riboswitches, the cis-regulatory elements present in the 5′ untranslated region of mRNA, that control gene expression by undergoing conformational transitions.^1,2^ Riboswitches generally consist of two domains: an aptamer domain (AD) and an expression platform (EP). The riboswitch cognate ligands bind to the AD, triggering an allosteric structural change in the EP that regulates transcription or translation events.^1–3^ A generic feature of signal transduction from the AD to the EP involves reorganization of hydrogen-bond networks.^2,4–6^ In riboswitches under transcriptional control, structural reorganization occurs between the anti-terminator and terminator stems in the EP. Similarly, in riboswitches under the translation control, the structural organization leads to the anti-sequestration or sequestration of the Shine-Dalgarno (SD) sequence where the ribosome binds to initiate translation.^2^ High-resolution three-dimensional structures of the riboswitches with both the AD and EP containing both ON and OFF states are rare. Therefore, investigating the molecular mechanism of signal transduction from the *apo* to the *holo* state is difficult, and the understanding is limited. ^7^

In this context, the S-adenosyl methionine (SAM) (Fig. 1a) sensing translational riboswitch, SAM-III, is an excellent model system for investigating the mechanism of signal transduction between the AD and EP,^8^ as high resolution crystal structures of the *apo* (ON) (PDB ID: 6C27)^9^ and *holo* (OFF) (PDB ID: 3E5C)^8^ states are available (Fig. 1b,c). SAM-III is an architectural outlier^8^ where the traditional modular boundary between the AD and EP does not exist, and the SD sequence is topologically integrated directly into the ligand-binding pocket (Fig. 1b,c). The *apo* structure consists of three helices (P0, P3, and P5) and a three-way junction (3WJ) that connects P0, P3, and P5 (Fig. 1b).^9^ In contrast, the SAM-bound *holo* structure consists of four helices (P1-P4), and a 3WJ connecting P1, P2, and P4 (Fig. 1c).^8^ The ligand-binding pocket is located in the 3WJ of the *holo*-state (Fig. 1c). Except for the P3 helix, the secondary and tertiary structures of the *apo* and *holo* states are different. To crystallize the *apo* state, the residues C48-U60 were deleted to destabilize the P1 and P4 helices (Fig. 1b,c). Conversely, to obtain the *holo* crystal structure, residues G1-A7 were removed to destabilize the competitive P0 helix (Fig. 1b,c). The SAM-binding pocket is absent in the *apo* state, raising the question of how SAM is recognized in the *apo* state.^4,9^

**Figure 1:**
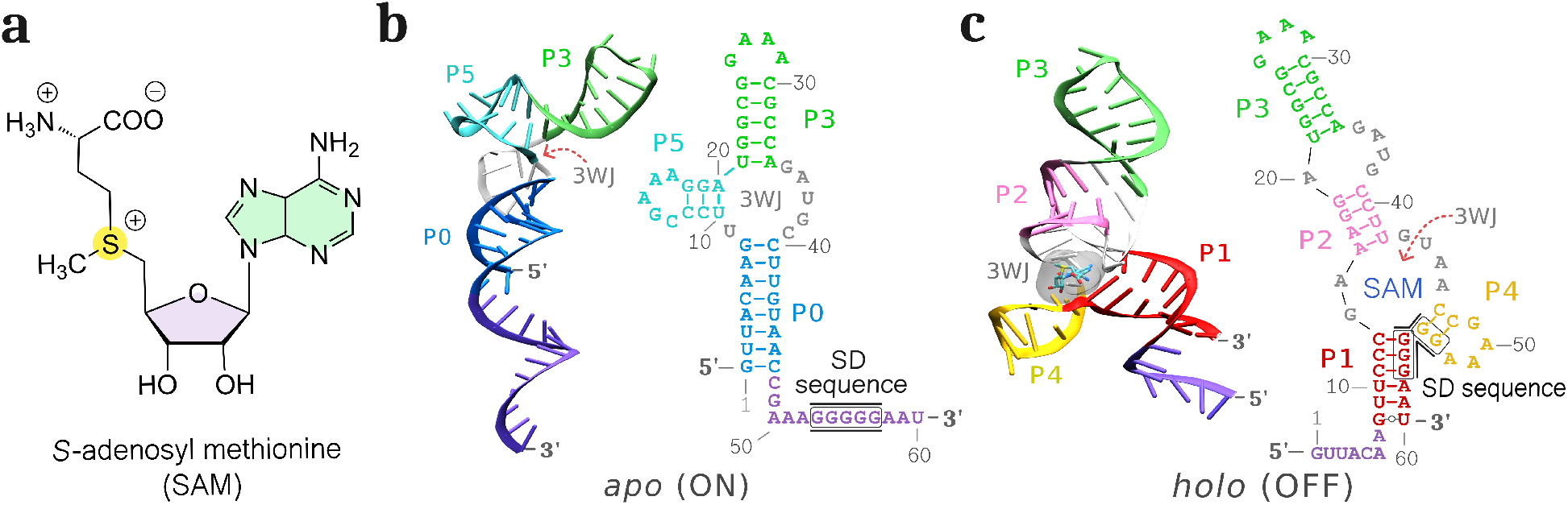
**(a)** Chemical structure of S-adenosyl methionine (SAM). Crystal structure and secondary structure of SAM-III riboswitch in **(b)** *apo* state (PDB ID: 6C27),^9^ and **(c)** *holo* state (PDB ID: 3E5C).^8^ Nucleotides added to the *apo* (C48–U60) and *holo* (G1–A7) crystal structures to make the sequence lengths identical are shown in violet.

Several experiments, including nuclear magnetic resonance (NMR), small-angle X-ray scattering (SAXS), isothermal calorimetry (ITC), and SHAPE analysis, have been performed to investigate ligand recognition.^4,5^ The NMR and SAXS experiments^4^ support the existence of a distinct *apo* structure consisting of P0 and P3 helices but did not comment on the presence of the P5 helix, as observed in the crystal structure.^9^ NMR and SAXS experiments^5^ further suggest the existence of a pre-bound or READY (R) state that connects the *apo* and *holo* states in equilibrium, 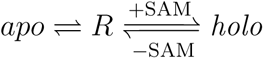. The structure of the R state is suggested to resemble that of the *holo* state, but the binding pocket remains open. The ITC experiments showed that, in the absence of SAM, *apo* remains as the major conformation (∼ 80%), and R remains as the minor conformation (∼ 20%).^4^ In the presence of SAM, the R state binds SAM, shifting the equilibrium to the right and forming the *holo* state. The SHAPE analysis also supported the existence of the R state.^5^

Despite these studies, crucial questions remain unclear – (i) What is the structure of the R state? (ii) How does the SAM-III riboswitch reorganize its hydrogen bond network from the *apo* to the R state transition? (iii) Are intermediate states populated in the transition path, and is the transition path unique? (iv) What role do metal ions play in governing the conformational pathways? In the experiments, it is extremely difficult to capture transient intermediate states. The use of all-atom simulations to investigate the transition landscape between the *apo* and R states is also challenging, as these limitations stem from slow transition kinetics and the complex, rugged free-energy landscape of RNA. To overcome these challenges, we performed molecular dynamics simulations using the three-interaction-site (TIS) coarse-grained (CG) RNA model, which was successfully used to investigate the folding thermodynamics and kinetics for diverse RNA systems. ^10–16^

We show by computing the free-energy landscape that *apo* conformation of SAM-III exists in a dynamic, ‘breathing-like’ equilibrium – fluctuating between open and closed states of its 3WJ domain, a process significantly affected by the Mg^2+^ concentration ([Mg^2+^]). We further identified the tertiary structure of the R state, and identified the critical intermediate states populated during the transition between the *apo* and R states. Due to intermediates, the transition route between the *apo* and R state is not unique and proceeds through multiple pathways. Interestingly, the preferred pathway for *apo* to R transition is highly dependent on [Mg^2+^]; however, the reverse transition is independent of [Mg^2+^]. Overall, this work reconciles previous experimental and theoretical findings and presents a comprehensive transition landscape of the SAM-III riboswitch, with implications for controlling gene regulatory pathways^4,5,17,18^ and effective biosensor engineering.^19,20^

## Results

### Simulations Capture Thermodynamic and Structural Features of the Ligand-free SAM-III Ensemble

The conformational switching between the *apo* (ON) and *holo* (OFF) states involves significant structural rearrangement; consequently, the emergence of intermediate states along the switching pathway is highly probable. SAXS, NMR, and ITC experiments collectively suggest that in the absence of SAM ligand, the SAM-III riboswitch mostly exists in the *apo* state with a low population of *holo*-like READY (R) state that possesses a pre-formed SAM-binding pocket, which readily binds to SAM and transits to the *holo* state.

To study the mechanism of *apo* ⇌ *holo* conformational switching in the SAM-III riboswitch, we performed simulations using the CG TIS RNA model and dual-basin potential energy surface constructed by combining the native contact information in the *apo* and *holo* crystal structures^8,9^ (PDB ID: 6C27 and 3E5C) (see Methods). We performed simulations of full 60-nt unfolded sequence (G1-U60) of SAM-III in the absence of SAM at four different temperatures, *T*_sim_ = 295, 310, 338 and 368 K, with [Mg^2+^] = 2, 5, and 8 mM, with a fixed background [K^+^] = 150 mM (Table S1).

To identify different thermodynamic states and their relative stabilities we calculated the one-dimensional free-energy-surface (1D FES) projected onto the difference between the *holo*-specific and *apo*-specific fraction of native contacts 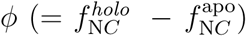 (see Data Analysis in Methods) (Fig. 2a). The conformations with *ϕ <* 0 are similar to *apo* state, and *ϕ >* 0 are similar to *holo* state. The 1D FES shows two adjacent states A1 (*ϕ* = −0.45) and A2 (*ϕ* = −0.3) representing the *apo* ensemble. The broad minima around *ϕ* ≈ 0.25 represent the holo-like R state, and a few intermediate states exist between *ϕ* = −0.15 and *ϕ* = 0.15 (Fig. 2a).

**Figure 2:**
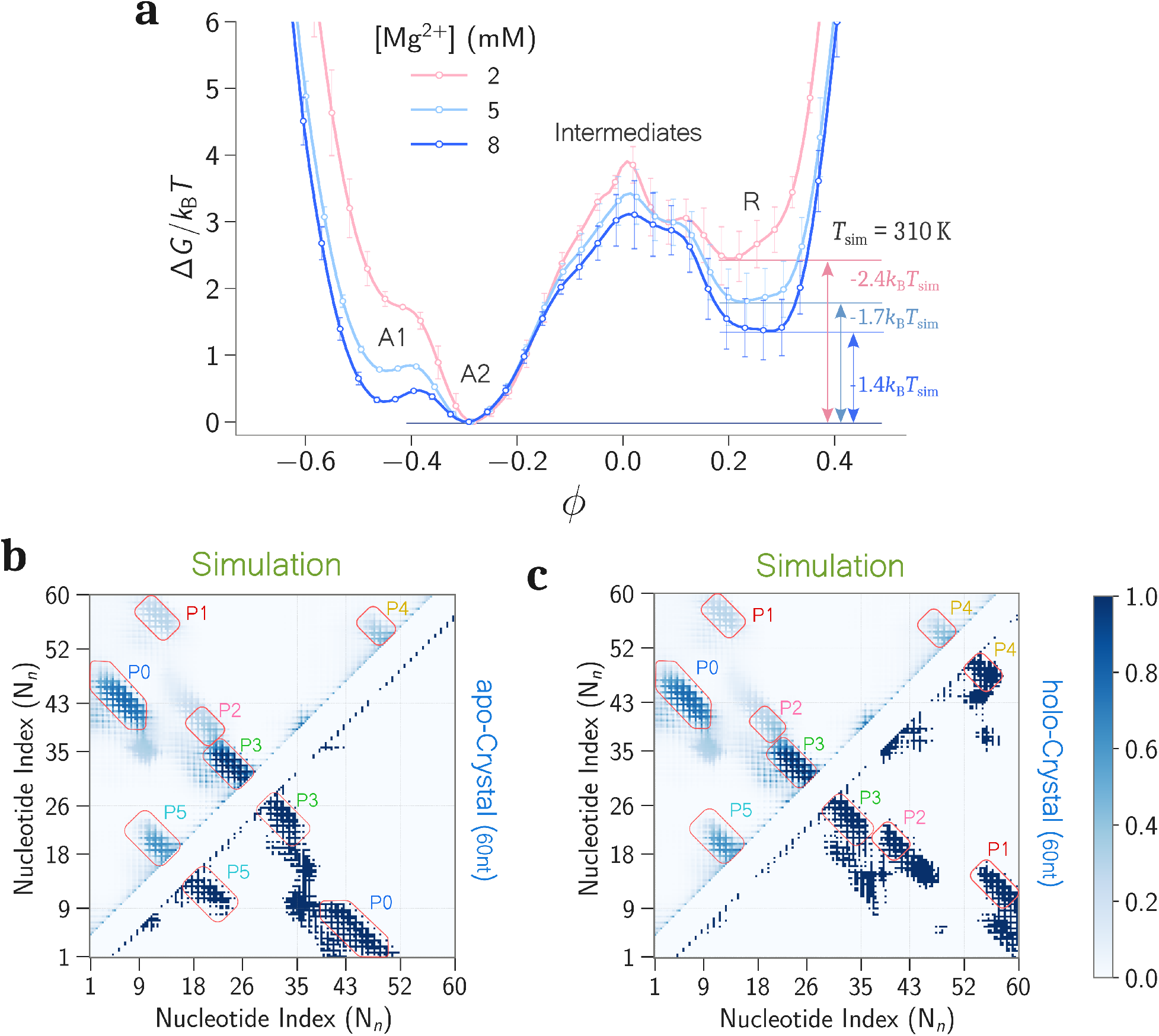
**(a)** 1D FES projected onto the difference between the fraction of native contacts between the *apo* and *holo* crystal structures, 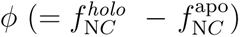 at 310 K, [K^+^] = 150 mM, and [Mg^2+^] = 2 mM, 5 mM and 8 mM. Comparison of the average contact map of SAM-III conformational ensemble obtained at *T*_sim_ = 310 K and [Mg^2+^] = 5 mM with the **(b)** *apo* and **(c)** *holo* crystal structures. The contact maps in the upper and lower diagonals are from simulations and the crystal structure, respectively. In the conformational ensemble obtained from the simulations, helices P0 and P3 are most stable.

The 1D FES clearly suggests that the apo ⇌ R transition is a multi-step process. We observed that the free-energy difference 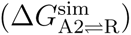 between the A2 and R states is ≈ 1.7 *k*_B_*T* (≈ 1.05 kcal mol*^−^*^1^) at [Mg^2+^] = 5 mM and *T*_sim_ = 310 K, which is in agreement with the ITC experiments^4^ 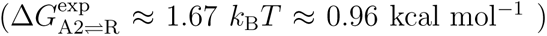 at [Mg^2+^] = 5 mM and *T*_exp_ = 288 K (see Data Analysis in SI). Thus, the ensemble sampled at *T*_sim_ = 310 K provides a thermodynamically calibrated representation of the experimentally observed ensemble at *T*_exp_ = 288 K. The difference between *T*_sim_ and *T*_exp_ arises from the simplified nature of the coarse-grained RNA model. In this model, the simulation temperature serves as the only adjustable parameter and was chosen to match the experimental free-energy scale.

To probe the structural features of conformations, we computed the average contact map of all the conformations sampled at *T*_sim_ = 310 K and [Mg^2+^] = 5 mM, and compared it with the *apo* and *holo* crystal structures (Fig. 2b,c). The contact map shows that in the absence of SAM binding, helices P0 and P3 are stable. In contrast, helix P5 is partially stable, and helices P1, P2, and P4 are unstable (Fig. 2b,c). This indicates that the overall conformational ensemble of SAM-III in the absence of SAM binding is dominated by *apo*-like structures (Fig. 1b). The simulation results are in agreement with the NMR study^4^ on SAM-III(59) (G1–A59), which showed dominant signals only for helices P0 and P3 at *T*_exp_ = 288 K.

We computed the pair-distance distribution (PDD) at different temperatures and compared it with the SAXS experiments^4^ on SAM-III(59) in the absence of SAM. We observed that the PDD obtained from *T*_sim_ = 368 K is quantitatively closest to that from SAXS at *T*_exp_ = 295 K with [Mg^2+^] = 5 mM (Fig. S1). The average radius of gyration from the simulation, 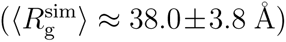 is close to the value reported from the SAXS data^4^ 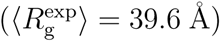, which indicates the dominance of the extended structures in the conformational ensemble. Based on this SAXS study, it was inferred that the broad PDD was due to structures in which P3 and P0 were connected via single-stranded regions. However, the average contact map computed at *T*_sim_ = 368 K shows a partially stable P3 helix, and the rest of the helices are unstable (Fig. S2).

### Mg^2+^ Modulates the ON/OFF Switching Landscape of SAM-III

The 1D FES clearly shows the shift in equilibrium from A2 to A1 with increasing [Mg^2+^] from 2 to 8 mM (Fig. 2a). However, A2 remains the global minima at [Mg^2+^] = 8 mM at *T*_sim_ = 310 K. This suggests that the equilibrium within the *apo* sub-states (A1 and A2) is strongly influenced by the Mg^2+^ ions. The 1D FES further shows that the broad R state also becomes more stable and with increasing [Mg^2+^] from 2 to 8 mM as indicated by the shifting of the minima from *ϕ* = 0.2 to *ϕ* = 0.3 (Fig. 2a). The average contact maps for the conformations around *ϕ* = 0.2 to *ϕ* = 0.3 in the R state show that the P4 helix is absent near *ϕ* = 0.2 and present near *ϕ* = 0.3 (Fig. S3a). This suggests that increasing [Mg^2+^] stabilizes the P4 helix, thereby promoting formation of the SAM-binding pocket (Fig. 2a).

The 1D FES further shows that as [Mg^2+^] is increased from 2 to 8 mM, the free-energy difference between the A2 and R states decreases from 2.4 to 1.4 *k*_B_*T* (*T*_sim_ = 310 K), suggesting that Mg^2+^ stabilizes the R state (Fig. 2a). However, it is important to note that even at high [Mg^2+^] (= 8 mM), the *apo*-like states (A1 and A2) remain the global minima, indicating that within the physiological Mg^2+^ concentration range, the R states are not the global minimum relative to the *apo*-like states (Fig. 2a).

To resolve the intermediate states populated during the transitions between the *apo* and R states we computed the two-dimensional (2D) FES projected onto the fraction of native contacts with respect to the *apo* 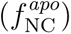 and *holo* 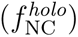 crystal structures at three different temperatures (Fig. 3, S4, S5 and S6) (see Data Analysis in Methods). At *T*_sim_ = 295 K, the 2D FES clearly shows five distinct conformational states, labeled A1, A2, I1, I2, and R (Fig. 3a). At this temperature, the transition between the *apo* and *holo*-like structures involves substantial structural rearrangement in the hydrogen-bonding network, which takes place through the population of I1 and I2 intermediate states.

**Figure 3:**
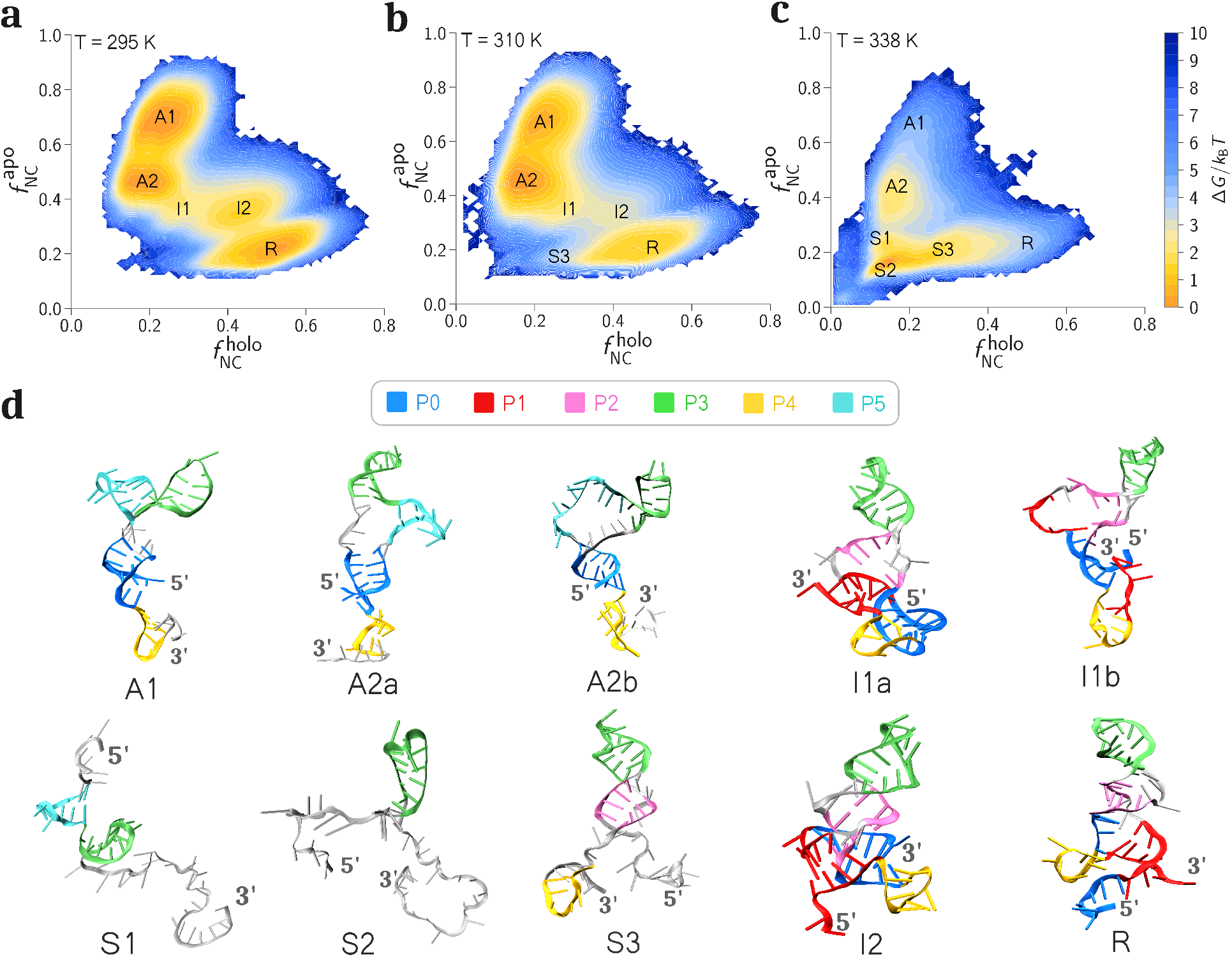
2D FES projected onto 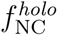 and 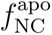 at [Mg^2+^] = 8 mM, [K^+^] = 150 mM, and **(a)***T*_sim_ = 295 K, **(b)** *T*_sim_ = 310 K, and **(c)** *T*_sim_ = 338 K. **(d)** Representative tertiary structures from different states. The basins A2 and I1 contain two sub-ensembles of conformations (A2a, A2b, I1a, I1b). The color scheme for different secondary structural elements is shown in the figure.

**Figure 4:**
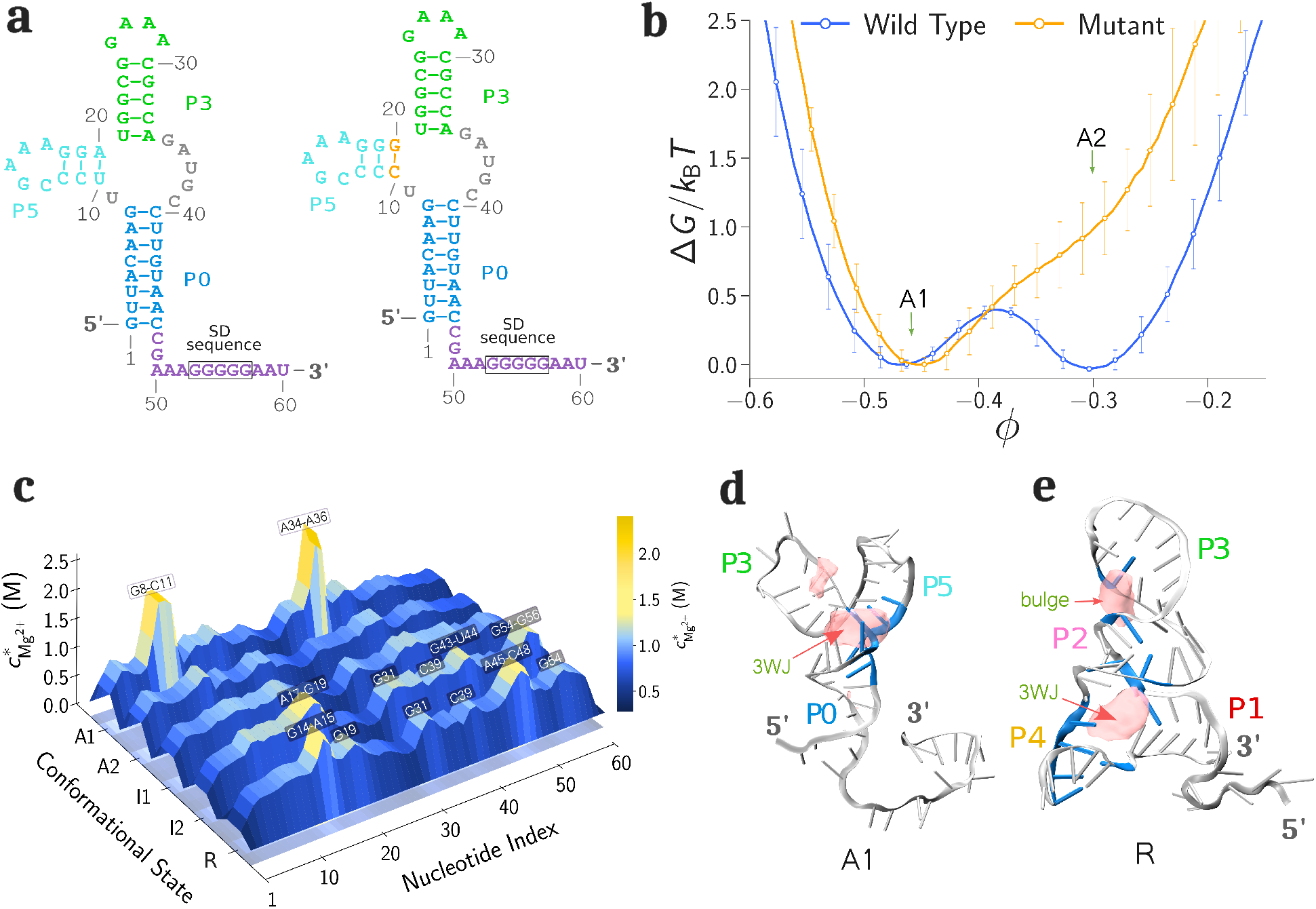
**(a)** The *apo* secondary structure in wild type and mutated sequences. The mutations in the P5 helix are U10C and A20G shown in orange. **(b)** 1D FES projected onto the difference between the *apo*-specific and *holo*-specific fraction of native contacts 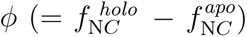 for wild type and mutated P5 at [Mg^2+^] = 5 mM, [K^+^] = 150 mM and 295 K. **(c)** 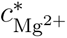 profile for the five states shows critical peaks for A1, I2 and R states at [Mg^2+^] = 8 mM. Spatial density map of Mg^2+^ ions around **(d)** A1 and **(e)** R states are shown as mauve iso-surfaces corresponding to the ISO value of 0.005, highlighting the high Mg^2+^ condensed region.

**Figure 5:**
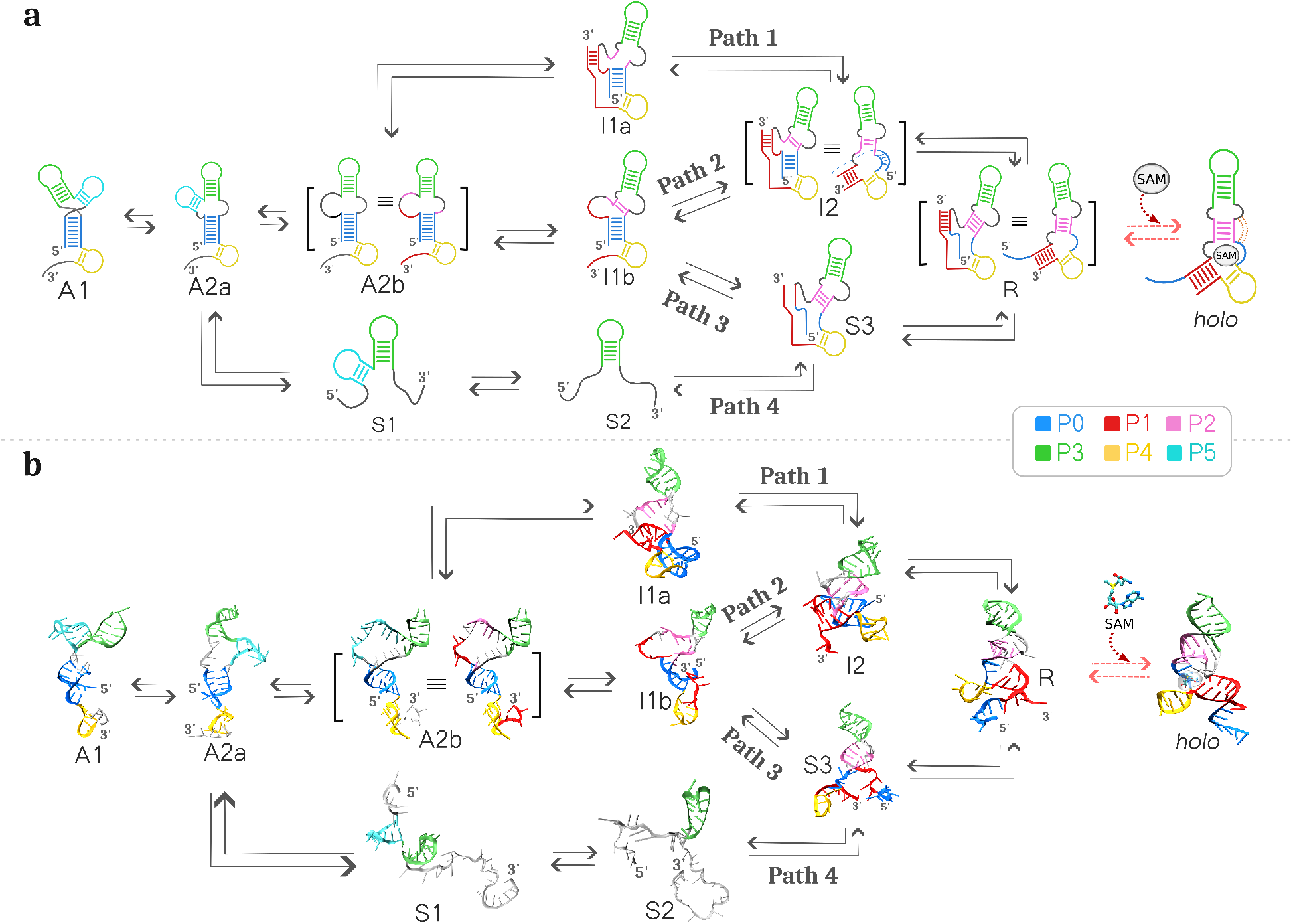
Transition paths associated with *apo* ⇌ R transformation at [Mg^2+^] = 8 mM and 310 K. **(a)** Schematic representation of transitions between identified conformational states. **(b)** Tertiary structures in transition pathways from the simulations. The final transition between R and *holo* is already known and depicted as red dashed arrows only for completeness of the transition landscape and is not a part of our simulations.^4,5,21,23^

At *T*_sim_ = 310 K, the I1 state became less stable and its basin merged with the A2 basin (Fig. 3b). The stability of the I2 state also decreased (Fig. 3b). However, the major states, *apo* (A1 and A2) and R, remain stable at this temperature. At *T*_sim_ = 338 K, we observed substantial changes in the 2D FES (Fig. 3c). Both the *apo* states, A1 and A2, and the R state are destabilized, and new low-free-energy regions corresponding to semi-folded states labeled S1, S2, and S3 are populated (Fig. 3c).

### *Apo*-state Exists in a Two-state ‘Breathing-like’ Dynamic Equilibrium

Conformations with 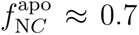 and 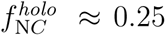 referred to as state A1 closely resemble the native *apo* state (Fig. 3a). The average native contact map of conformations in the A1 basin confirms that the helices P0, P3, and P5 in these conformations are stable along with the 3WJ, confirming a high degree of structural resemblance with the *apo* crystal structure^9^ (Fig. S7). Conformations with 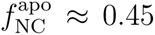 and 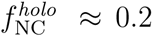 referred to as state A2 also resembles the native *apo*-state (Fig. 3a). The average native contact map of conformations in the A2 basin shows that the helices P0 and P3 helices are stable, helix P5 is partially formed, and the contacts associated with the 3WJ are entirely missing (Fig. S8). We further observed that the conformational transitions between the A1 and A2 states are rapid, indicating a fast ‘breathing-like’ motion of the 3WJ, in which the 3WJ opens in A2 and closes in A1 (Fig. S9).

To test the effect of helix P5 on the breathing of the 3WJ in the *apo* conformations, we introduced two stabilizing mutations in the P5 helix (U10C and A20G) at 3WJ, replacing an AU with a GC basepair (Fig. 4a). We performed CG simulations of this mutant sequence at [Mg^2+^] = 5 mM, [K^+^] = 150 mM and *T*_sim_ = 295 K, starting from the native *apo* structure (Fig. 1b). Comparison of the 1D FES of the wild type and mutant sequence projected onto 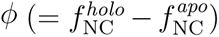 shows that the *apo* population is shifted towards A1 in the mutant sequence (Fig. 4b). This shift shows that P5 stabilization favors a more compact and closed 3WJ, thereby suppressing its ‘breathing-like’ motions (Fig. S10). Consequently, the increased A1 population suggests that, in addition to the P0 stability, P5 stability is also crucial for maintaining the structural integrity of the *apo* structure and can influence transitions to the R state.^4^

The ensemble of conformations in the A2 state is further divided into two sub-ensembles: one with a stable P5 helix (A2a) and the other with an unfolded P5 helix (A2b). We computed the 1D FES of the conformations in the A2 basin by projecting it onto the fraction of P5 native contacts 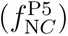, which clearly shows the A2a and A2b sub-ensembles, and the relative population of A2a is slightly dominant over A2b (Fig. S11, S12 and S13). The native contact maps of conformations in the A1 and A2 states also reveal the incipient formation of the P4 helix, which is a feature of the *holo* structure (Fig. S7a, S8). This premature P4 formation in an *apo*-like state (A1, A2) stems from the sequence positioning of P4 nucleotides at the 3′-terminus following the P0 helix (Fig. 1b). This arrangement effectively isolates P4 from the core *apo* architecture (P0, P3, P5, and the 3WJ), allowing it to fold independently.

### Intermediates Have Hybrid Structural Features from *apo* and R States

The average native contact map of the conformations in the state I1 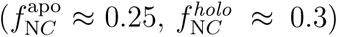 shows that conformations have structural components from both *apo* and *holo* crystal structures (Fig. S14). The helices P0, P1, and P3 are prominently present, but P2 is partially formed in this state. There are two sub-ensembles of conformations populated in the I1 basin, and to classify them, we projected the conformations in the I1 basin onto a 2D FES with 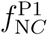 and 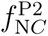 as the CVs (Fig. S15). In the sub-ensemble I1a, the helices P0, P1, and P3 are present (Fig. S16), and P2 is absent. In the sub-ensemble I1b, the helices P0, P2, and P3 are present, and P1 is absent (Fig. S17). The sub-ensemble I1a is relatively more stable than I1b. The conformations in the state I2 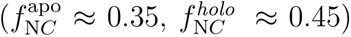 consist mainly of the common P3 helix, helical components P1, P2, and P4 from the *holo* structure and a significant helical component P0 from the *apo* structure (Fig. S18).

### Native Contacts in the SAM-binding Pocket Are Not Stable in the Absence of SAM

The R state conformational basin is broad with a minima at 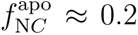 and 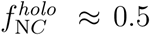 (Fig. 3a-b). The average native contact map of conformations in the R basin shows that the P1, P2, P3 helices in the *holo* structure were formed, and the P4 helix is partially formed (Fig. S19). The majority of contacts in the SAM-binding pocket (BP1, BP3, BP4, BP5), except for BP2 are poorly formed (Fig. S19). This indicates that in the absence of cognate ligand binding, the SAM-binding pocket loses its compactness. Consequently, the SAM-III riboswitch does not remain stable in its *holo* crystal geometry and populates the R state. The NMR^4^ and SHAPE^5^ experiments designed to probe the R state suggest the existence of a flexible and open binding pocket in the R state. The atomistic simulations performed on the 53-nt *holo* crystal structure^8^ also suggested the existence of the dynamic and weakly structured SAM-binding pocket in the binding competent state in the absence of SAM.^21–23^ Therefore, the R state observed in our simulations is in agreement with the previous experimental and theoretical studies.

### Specific Mg^2+^ Condensation Is Critical for the Stabilization of *apo* States

We computed the local Mg^2+^ concentration 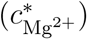 around the phosphate backbone (Eq. (2)) of the riboswitch for all the states populated in the simulations at [Mg^2+^] = 8 mM and *T*_sim_ = 310 K to identify the preference for Mg^2+^ to condense in specific regions of their structures (Fig. 4c, S20). For the state A1, the 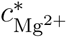 profile shows two sharp spikes near G8-C11 and A34-A36 indicating specific Mg^2+^ condensation at the 3WJ confirming that Mg^2+^ affects the breathing-like dynamics of the 3WJ in state A1. The spatial density map of Mg^2+^ also confirms its condensation at the 3WJ (Fig. 4d) (see Data Analysis in SI). The strong condensation of Mg^2+^ ions in the 3WJ of state A1 (Fig. 4c) compared to any other state indicates that Mg^2+^ is critical to the stabilization of the native-like *apo* state similar to the crystal structure^9^ (Fig. 1b).

We showed that stability of the R state increased with [Mg^2+^] (Fig. 2a). The average native contact map for the R state showed that a substantial number of contacts associated with the SAM-binding pocket are unstable even at high [Mg^2+^] (= 8 mM) (Fig. S19a). The 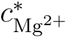 profile of R state shows specific spikes around G14-A15, G19, G31, A45-C48, and G54, indicating specific Mg^2+^ condensation at the 3WJ of the R state (Fig. 4c). Specifically, the spikes A45-C48 and G54 confirm the Mg^2+^-induced stability of the P4 in R state observed in 1D FES (Fig. 2a). However, the 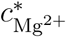 peaks are not large compared to A1. This suggests that Mg^2+^ alone cannot stabilize the majority of the contacts in the SAM-binding pocket to form the *holo* state. Thus, Mg^2+^ weakly stabilizes P4 and a few binding pocket contacts in the R state, thereby assisting the SAM-binding process. The spatial density map of Mg^2+^ also shows specific Mg^2+^ condensation at the 3WJ in R state (Fig. 4e). A weak Mg^2+^ condensation was also observed in the bulged region between the P2 and P3 helices.

The 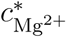 profile of I2 also shows peaks for specific nucleotides (Fig. 4c). The peaks at A17-G19 indicate condensation of Mg^2+^ at the binding pocket region, and the peaks at G43-U44 and G54-G56 indicate the condensation of Mg^2+^ to stabilize the partially formed P1 and P0 helices in I2. The states A2 and I1 show no significant peaks, indicating no specific Mg^2+^ binding (Fig. 4c).

### Transition Flux Between *apo* and R States is Distributed Among Multiple Switching Pathways

The *apo* → R transition requires extensive reorganization of the hydrogen-bond network and therefore proceeds through multiple intermediate states. Analysis of transition trajectories for [Mg^2+^] = 2, 5, and 8 mM at *T*_sim_ = 310 K reveals a branched network of pathways connecting the *apo* and R states (Fig. 5). We identified four major transition pathways from this network (Table 1). **Path 1:** Starting from A1, the riboswitch populates A2a through opening of the 3WJ, followed by A2b through disruption of P5 and partial destabilization of P0 near the junction (Fig. 5). From A2b, one of the freely moving 3′-terminal strands (G55–U60) partially pairs with its complementary strand (G8–C13) in the open 3WJ of A2b to form P1, populating I1a. Partial formation of P2 then leads to I2, followed by the complete disruption of the remaining P0 helix to yield the R state (Fig. S21). **Path 2:** This pathway is identical to Path 1 up to the formation of A2b (A1→A2a→A2b), and then proceeds through I1b, where P2 forms partially before P1. Subsequent formation of P1 converts I1b to I2, which then progresses to form R by breaking the P0 (Fig. S22). **Path 3:** This route is identical to Path 2 up to the formation of I1b (A1→A2a→A2b→I1b). From I1b, complete disruption of P0 gives rise to the semi-folded intermediate S3, from which P1 formation leads to the R state (Fig. S23). A helix-based kinetic modeling study^18^ of SAM-III predicted that the transition proceeds via a single route, which is similar to the Path 3 observed here (Fig. 5). **Path 4:** This pathway proceeds mainly through semi-folded intermediates. Initially, there is a transition from A1 to A2a, then P0 disrupts to form S1, followed by the disruption of P5 to yield the extended intermediate S2, in which only P3 is stable. Formation of P2 converts state S2 to S3, followed by the P1 formation to yield the R state (Fig. S24).

**Table 1:**
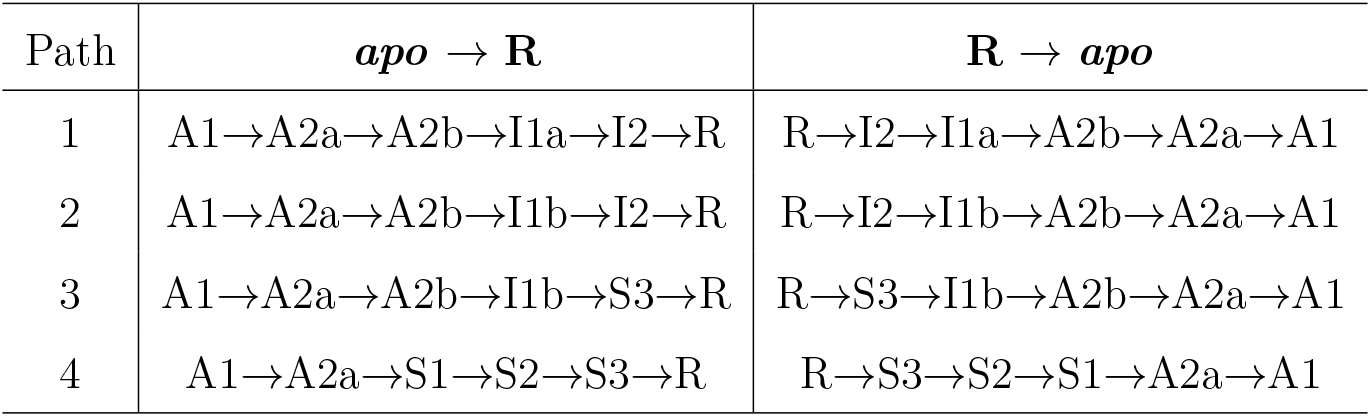
Major Transition Pathways.

| Path | *apo* → **R** | **R** → *apo* |
| --- | --- | --- |
| 1 | A1→A2a→A2b→I1a→I2→R | R→I2→I1a→A2b→A2a→A1 |
| 2 | A1→A2a→A2b→I1b→I2→R | R→I2→I1b→A2b→A2a→A1 |
| 3 | A1→A2a→A2b→I1b→S3→R | R→S3→I1b→A2b→A2a→A1 |
| 4 | A1→A2a→S1→S2→S3→R | R→S3→S2→S1→A2a→A1 |

The trajectories show that reverse, R → *apo*, transition proceeds through the same four pathways in the opposite direction (Fig. S25 to S28). Previous experiments^24^ and theoretical studies^18,25^ also indicated that the SAM-III riboswitch operates as a reversible switch, corroborating our observation of the reversible switching pathways.

### Mg^2+^ Asymmetrically Influences *apo* → R and R → *apo* Transition Pathways

We computed the probabilities of selecting the transition pathways between the *apo* and R states at *T*_sim_ = 310 K for [Mg^2+^] = 2, 5, and 8 mM (see Data Analysis in SI). We observed that the probabilities of the A1 → R transition pathways strongly depend on [Mg^2+^] (Fig. 6a). At [Mg^2+^] = 2 mM, the flux is distributed across multiple routes. The probabilities of the paths follow the order: Path 3 (0.37) *>* Path 1 (0.27) *>* Path 2 (0.19) *>* Path 4 (0.17). At [Mg^2+^] = 5 mM, the flux strongly shifts toward Path 1, and the transition path flux follows the order: Path 1 (0.53) *>* Path 3 (0.23) *>* Path 2 (0.17) *>* Path 4 (0.07). At [Mg^2+^] = 8 mM, transition through Path 1 dominates and follows the order: Path 1 (0.60) *>* Path 3 (0.21) *>* Path 2 (0.13) *>* Path 4 (0.06). Thus, with increasing Mg^2+^, a dominant transition path for A1 → R (Path 1) emerged, suppressing the alternate pathways (Fig. 6a).

**Figure 6:**
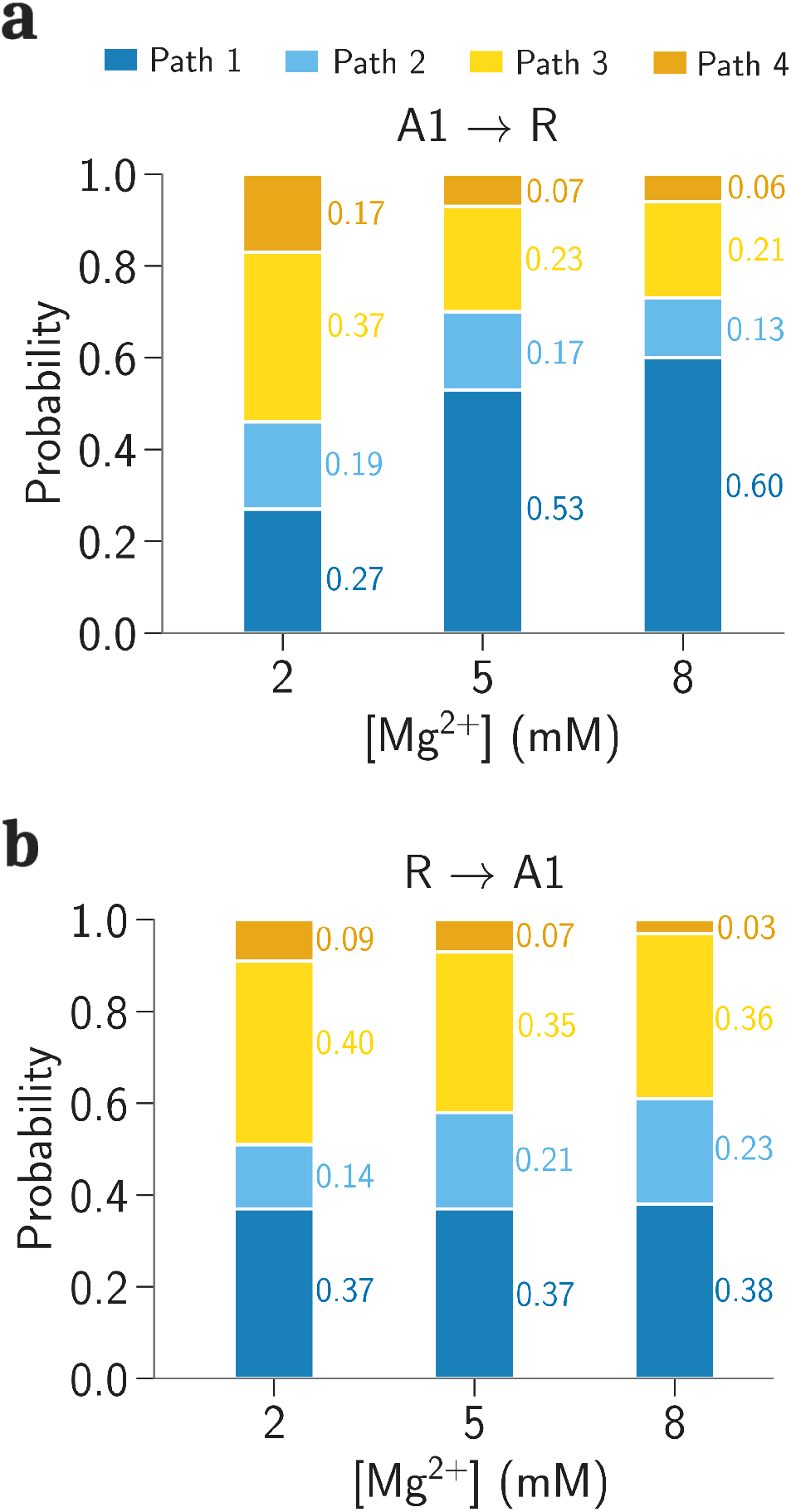
Probabilities of the various transition paths for **(a)** A1 →− *T*_sim_ = 310 K, [K^+^] = 150 mM, and [Mg^2+^] = 2, 5, and 8 mM.

In contrast, the R→A1 transition exhibits a weaker Mg^2+^ dependence (Fig. 6b). At [Mg^2+^] = 2 mM, the reverse transition mainly proceeds through Path 3 and follows the order: Path 3 (0.40) *>* Path 1 (0.37) *>* Path 2 (0.14) *>* Path 4 (0.09). At [Mg^2+^] = 5 mM, the transition is mainly through Path 1 and Path 3 and follows the order: Path 1 (0.37) *>* Path 3 (0.35) *>* Path 2 (0.21) *>* Path 4 (0.07). At [Mg^2+^] = 8 mM, the transition preference also follows a similar order: Path 1 (0.38) *>* Path 3 (0.36) *>* Path 2 (0.23) *>* Path 4 (0.03). Therefore, the R → A1 transition remains distributed primarily between Path 1 and Path 3 almost invariably across the entire Mg^2+^ range (Fig. 6b).

The difference in A1→R and R→A1 pathway preferences indicates that Mg^2+^ asymmetrically influences the forward and reverse transition landscapes. In the A1→R direction, the larger P0 helix should unfold, and increasing the [Mg^2+^] further inhibits this unfolding. Whereas increasing [Mg^2+^] facilitates the stepwise formation of the smaller P1 and P2 helices, and it further stabilizes compact intermediates such as I1a and I2 (Fig. 5). Hence, increasing [Mg^2+^] lowers the effective free-energy cost of progressing through Path 1 and concentrates the forward flux along this route. In contrast, the reverse transition requires formation of the larger P0 helix on top of the R state in which relatively smaller P1/P2 helices are present. This reorganization (R→I2) likely incurs a greater configurational penalty and is therefore less sensitive to the change in [Mg^2+^]. As a result, Mg^2+^ strongly channels the A1→R transition through Path 1 but has a weaker effect on the R→A1 transition, which remains distributed over multiple routes.

### Elevated Temperature Eliminates Transition Pathways with Compact Intermediates

To examine the effect of temperature on the probabilities of transition pathway selection, we compared the transition pathway selection probabilities at [Mg^2+^] = 8 mM, and *T*_sim_ = 310 K and 338 K. At 338 K, the *apo*→R and R→*apo* transitions mainly proceeds through paths 3 and 4, where compact intermediates I1 and I2 are not populated, and proceed through states with fewer native structural features. This redistribution of transition pathways with temperature is consistent with the FES, which shows that the native-like states (A1 and R) and compact intermediates (I1 and I2) lose stability, and the semi-folded states (S1, S2, and S3) with minimal native structure dominate (Fig. 3).

## Discussion

### SAM-III Conformational Ensemble Is Heterogeneous in the Absence of Ligand Binding

Our simulations reconcile the inferences from previous NMR, ITC, SAXS, ^4^ SHAPE,^5^ and X-ray crystallography^8,9^ experiments and provide a comprehensive mechanism of the ligand-free ON/OFF switching landscape of the SAM-III riboswitch. NMR and SAXS experiments^4^ infer that the full-length construct dominantly populates an *apo*-like state, which is similar to the A2 state, which we identified in our simulations (Fig. S12, S13). These experiments also indicate that the *apo* basin is internally broad: NMR^4^ detects a dominant *apo* ensemble defined primarily by P0 and P3, and SAXS^4^ reports a more extended ligand-free distribution than expected for a single compact *apo*-like structure, which corroborates our observation of the dynamic ‘breathing-like’ equilibrium between two *apo* states (A1 and A2) (Fig. 2a, 3a-d). NMR^4^ and SHAPE^5^ experiments further indicate that the binding pocket region in the ligand-free *holo*-like state remains more flexible than the SAM-bound *holo* (OFF) state. This agrees with our simulations, where the R state appears as a broad basin in the FES (Fig. 2a, 3a and 3b) with most of the binding-pocket contacts broken (BP1–BP5, except BP2)(Fig. S19). ITC experiments^4^ suggested the existence of an *apo*-dominated ensemble (*apo* ∼ 80% and R ∼ 20%) in the absence of SAM. This population balance is also approximately maintained in our ligand-free simulations (Fig. 2a). Thus, the dual-basin TIS RNA model captures the correct thermodynamic competition between *apo* and R states, while modestly favoring more compact microstates within the *apo* ensemble than are implied by SAXS and NMR studies.^4,5^

### SAM-III Follows a Sequential Conformational-Selection and Induced-Fit Mechanism for Ligand Binding

The results from the simulations in this study and the existing studies ^4,5,18,21,23,25^ support a ligand-recognition mechanism in the SAM-III riboswitch, best described as a sequential hybrid of conformational selection and induced fit. In the ligand-free ensemble, the riboswitch predominantly occupies an *apo*-like state but retains access to a smaller subpopulation of R that is structurally competent for SAM ligand binding, consistent with the conformational selection mechanism.^4^ After SAM binding, additional local reorganization of the binding-pocket and junctional regions is required to stabilize the fully repressed OFF state, consistent with the induced fit mechanism.^4,5,23^ This interpretation places the SAM-III riboswitch within a broader class of translational riboswitches that exploit pre-organized but still flexible *holo*-like states prior to ligand binding. The sequential combination of conformational selection and induced fit inferred here for the SAM-III riboswitch is consistent with mechanisms proposed for other metabolite-sensing riboswitches, including preQ1, lysine, guanine, adenine, and SAM-I riboswitches, where ligand binding occurs within a Mg^2+^ assisted pre-organized ligand-free ensemble but is followed by additional local ordering of the binding _core.7,26–31_

### Temperature, Mg^2+^, and Peripheral Helices Tune ON/OFF Switching Landscape in SAM-III

A major mechanistic insight from this work is that multiple intermediates are populated in the *apo* (ON state) → R (OFF state) transition path, and as a result, the switching transition proceeds through multiple competing routes rather than along a single deterministic pathway. At low temperatures, switching is biased towards proceeding via compact hybrid-helix intermediates, with gradual disruption of the P0 helix and the simultaneous formation of the P1/P2 helix. At elevated temperatures, the compact, native-like states and folded intermediates are destabilized, and the dominant switching pathway proceeds via semi-folded intermediates. In contrast, Mg^2+^ stabilizes the compact intermediates and increases flux through the direct helix-exchange route (Path 1) (Fig. 5, 6), where helical components from both *apo* and *holo* structures coexist. Thus, SAM-III behaves as a conformational switch, where the switching pathway is governed by the thermodynamic balance between compact and weakly structured states, tuned by [Mg^2+^] and temperature.

Beyond the effects of temperature and [Mg^2+^], the switching landscape is also expected to depend on stabilization of the ligand-binding core. However, the relative stability of peripheral helices P0 and P5, present in the *apo* state play an important role. The disruption of P0 helix biases SAM-III toward an irreversible R (OFF) state irrespective of ligand availability.^4,5^ Analogously, we showed using simulations that stabilizing the P5 helix through a base-pair mutation shifts the equilibrium further towards compact *apo* conformations (A2 ⇌ A1) (Fig. 4a,b), delaying premature access to the R (OFF) ensemble. Therefore, tuning the stability of P0 and P5 helices serves as a control for the SAM-III-based engineered biosensors.^19,20^

### Multiple ON/OFF Switching Routes Could be a General Feature of Translational SAM Riboswitches

Among translational SAM riboswitches, SAM-III closely resembles SAM-VI in sequence and structure, resulting in a similar switching architecture.^32,33^ Crystal structure of SAM-VI revealed a 3WJ with a short P3 helix and an SD sequence positioned between P1 and P3, similar to the structural features observed in SAM-III.^8,33^ Recent single-molecule and co-transcriptional studies further inferred that SAM-VI does not switch in a strictly two-state manner, but instead populates intermediates on the transition pathway, including mixed-helical conformations, before reaching the final SAM-bound repressed state.^33–35^ These intermediates are analogous to the SAM-III intermediates (I1 and I2), where P0, P1, and P2 helices coexist, observed in Paths 1 and 2 (Figure 5). However, our simulations show that SAM-III switching happens through similar folded intermediates predicted by the experiments^33–35^ on SAM-VI, and also the semifolded intermediates predicted by a theoretical study^18^ on SAM-III. Thus, we propose new experiments^36,37^ on SAM-III to test the existence of the compact intermediates such as I1 and I2, and experiments on SAM-VI to test the existence of semifolded intermediates (S1, S2, and S3) in their switching pathways to confirm if similar multiple switching routes are a general feature of translational SAM riboswitches.

## Conclusion

In summary, we provide a comprehensive mechanistic framework for reversible conformational switching in the SAM-III riboswitch for gene regulation. The ligand-free ensemble is dominated by an *apo*-like state but also populates a minor subensemble of READY (R) conformations that facilitate SAM ligand binding, rationalizing the conformational selection mechanism. The transition between the *apo*-like and R states proceeds through multiple structurally distinct intermediates and is strongly affected by [Mg^2+^] and temperature. Taken together with prior experimental studies, the present results support a hybrid mechanism for ligand binding by SAM-III involving conformational selection followed by ligand-induced local structural reorganization into a fully repressed OFF state. We further propose multiple new experiments to probe the structure of the intermediates populated during the conformational switching of the translational riboswitches, SAM-III and SAM-VI, to establish that multiple similar switching routes are a general feature of these riboswitches. A drawback of this study is that the dual-basin TIS RNA model relies on the native contacts from the *apo* and *holo* structures. Therefore, pathways primarily stabilized by non-native interactions may remain underrepresented. However, this is unlikely to affect the central conclusions, as the dominant switching routes identified here proceed through native-like intermediates that connect the experimentally supported *apo* and R ensembles and reconcile previous studies.^4,18,24,25,38^

## Methods

### Coarse-grained MD Simulations

Several CG RNA models have been developed to investigate structure-related problems in RNA systems.^7,13,14,39–41^ We used the TIS CG RNA model^10,39^ to investigate the conformational switching in the SAM-III riboswitch. In this model, each nucleotide is represented by three beads, which mimic the phosphate, sugar, and base groups. The solvent is treated implicitly through a temperature-dependent dielectric constant. K^+^ and Mg^2+^ ions are included explicitly to account for ion correlations and discrete RNA–ion interactions. The TIS model is described in detail in the supporting information (SI).

### Initial Structure Preparation for SAM-III Simulations

We prepared the initial structures for the simulations using the *apo* (ON) and *holo* (OFF) crystal structures (PDB IDs: 3E5C and 6C27, respectively).^8,9^ In the experiments, to crystallize the structures, portions of the full sequence were used along with the stabilizing mutations.^8,9^ The 47-nucleotide-long *apo* structure^9^ and the 53-nucleotide-long *holo* crystal structure^8^ share some common residues but also have some residues that are mutually exclusive. Therefore, we combined both the *apo* and *holo* sequences to obtain the full 60 nt sequence. In the apo crystal structure, the U2G, U3G, A45C, and A46C mutations were used for crystallization. However, we reverted them using COOT^42,43^ to keep the sequences identical in both *apo* and *holo* structures. We added residues C48-U60 at the 3′ end of the *apo* crystal structure and residues G1-A7 at the 5′ end of the *holo* crystal structure using COOT^42,43^ (Fig. 1b,c). This full sequence is identical to the sequence used by Wilson et al.,^4^ except one extra uracil (U) at the 3′-end, which is present in the *holo* crystal structure.^8^ We performed Langevin dynamics simulations using the extended-*holo* crystal structure (60 nt) (Fig. 1c) as the initial conformation to generate an ensemble of completely unfolded structures for subsequent simulations. These simulations were performed at a high temperature (373 K), and the hydrogen-bond potential (*U*_HB_) (Eq. S1) was disabled to unfold the structure quickly.

### Simulation Protocol

To investigate the mechanism of conformational transitions between the two states, we combined the native structural information from both the *apo* and *holo* crystal structures to construct a dual-basin framework^7,44^ within TIS force field. Using the previously generated unfolded structures of the 60 nt full construct as initial conformations, we subsequently performed multiple independent simulations at different temperatures and Mg^2+^ concentrations (Figure 1b,c, Table S1). We used a cubic box with an edge length of 200 ^°^A for all simulations. We implemented the Langevin dynamics simulations of the TIS RNA model in OpenMM.^45^ Further simulation details are provided in the SI.

### Data Analysis

#### Fraction of Native Contacts

The fraction of native contacts (*f_NC_*) between a particular set of nucleotides (*λ*, see Table S5) in a conformation *i*, is computed using 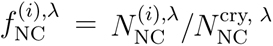, where 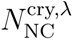 and 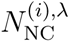 are the number of native contacts present between the nucleotides belonging to set *λ* in the corresponding *apo* or *holo* crystal structure and the *i*^th^ conformation, respectively.^46^ A pair of beads is considered to be in contact if the distance between them is less than the cutoff distance *r_cut_*(= 15 ^°^A).^10,47^

#### Free Energy Surface (FES) from CG Simulations

The FES projected onto the set of collective variables {*α*} is computed using the equation:

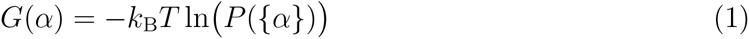

where *k*_B_ is the Boltzmann constant, *T* is the temperature, and *P* ({*α*}) is the joint probability distribution of the set of collective variables {*α*} at a given [Mg^2+^] and *T*. We computed the 1D FES calculations by projecting onto the collective variables 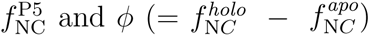. We computed 2D FES to characterize the various thermodynamic states populated during the structural transitions between the *apo* and *holo* structures by projecting onto the collective variables 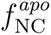 and 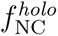. We also projected 2D FES to classify the I1 sub-ensembles using collective variables 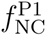 and 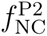.

#### Local Ion Concentration

The local Mg^2+^ concentration around a specific backbone phosphate bead in the RNA sequence is computed using, ^10,46^

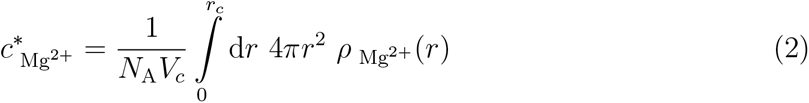

where 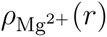 is the number density of Mg at a distance *r* from the backbone phosphate bead under consideration, *V_c_* is the spherical volume with a cutoff radius *r_c_* (= 5 ^°^A), and *N*_A_ is the Avogadro number.

## Supporting Information

Simulation descriptions and data analysis; description of CG simulation systems (Table S1); coarse-grained parameters (Table S2 to S4); description of nucleotides of sub-parts of SAM-III (Table S5); pair distance distribution of SAM-III(59) (Fig. S1); average contact maps of SAM-III (Fig. S2); average contact maps with schematic and tertiary structures of different states (Fig. S3, S7, S8, S12 to S14, S16 to S19); 2D FES at different [Mg^2+^] and temperatures (Fig. S4, S5); trajectories at different temperatures (Fig. S6,S9, S10); FES projected on 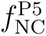 (Fig. S11); 2D FES projected on 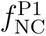 and 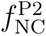 (Fig. S15); local 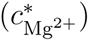 profile (Fig. S20); representative trajectory segments showing all transition pathways listed in Table 1 (Fig. S21 to S28).

## Acknowledgment

GR acknowledges funding from the Science and Engineering Research Board (SERB) through the grant CRG/2023/002817. DM acknowledges the research fellowship from the Indian Institute of Science, Bangalore. SPC acknowledges the INSPIRE research fellowship from the Department of Science and Technology (DST) India. We acknowledge the National Supercomputing Mission (NSM) for providing computing resources of “Param Pravega” at IISc supported by the Ministry of Electronics and Information Technology (MeitY) and the DST, Government of India. This research also used resources of the National Energy Research Scientific Computing Center (NERSC), a Department of Energy User Facility using NERSC award ERCAP0037272.

## For Table of Contents Use Only

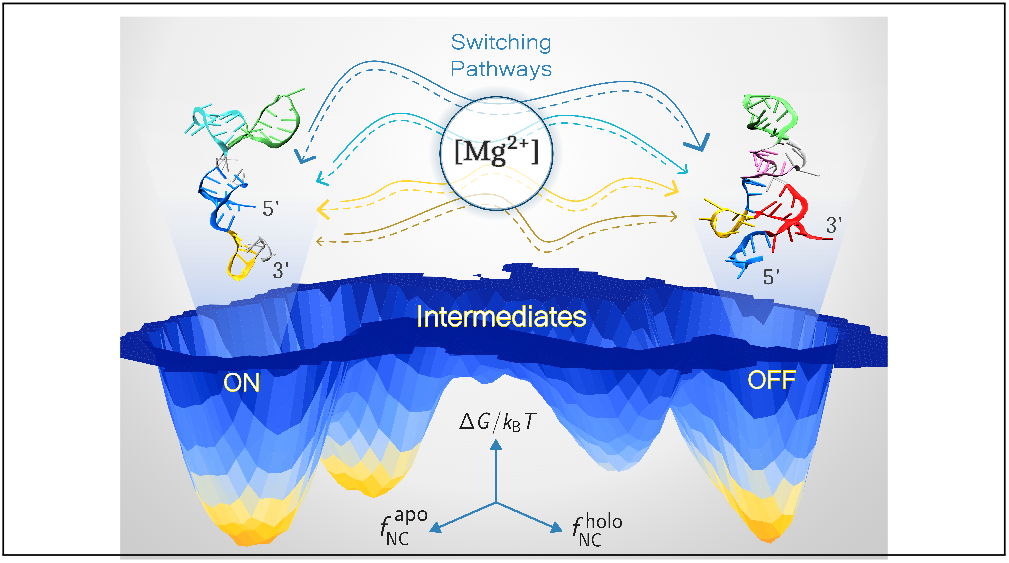

