## Supporting Information for "Conformational Switching between ON and OFF States Proceeds through Multiple Pathways in the SAM-III Translational Riboswitch"

### Simulation Descriptions

#### Three-Interaction-Site (TIS) RNA Coarse-Grained (CG) Model

The CG simulations were performed using the three-interaction-site (TIS) RNA model.[1,2] In this model, each nucleotide is represented by three beads centered on the phosphate group (P), sugar (S), and nucleobase (B), respectively. The TIS energy function[1] is given by

$$U_{\text{TIS}} = U_{\text{BL}} + U_{\text{BA}} + U_{\text{EV}} + U_{\text{EL}} + U_{\text{ST}} + U_{\text{HB}} + U_{\text{TST}} \tag{S1}$$

where the terms represent potentials for bond length ($U_{\text{BL}}$), bond angle ($U_{\text{BA}}$), excluded volume interactions ($U_{\text{EV}}$), electrostatic interactions ($U_{\text{EL}}$), stacking between consecutive bases ($U_{\text{ST}}$), tertiary stacking between non-consecutive bases ($U_{\text{TST}}$), and native hydrogen bond interactions present in the folded structure ($U_{\text{HB}}$).

The bond length and bond angle interactions are modeled using the harmonic potentials:

$$U_{\text{BL}} = k_\rho(\rho - \rho_0)^2 \tag{S2}$$

$$U_{\text{BA}} = k_\alpha(\alpha - \alpha_0)^2 \tag{S3}$$

where $\rho_0$ and $\alpha_0$ denote the equilibrium bond distance and bond angle, respectively, obtained from the ideal A-form RNA helix. The values of $k_\rho$ are 23, 64, and 10 kcal.mol$^{-1}$·Å$^{-2}$ for the P→S, S→P, and S−B bonds, respectively, where → indicates the $5' \rightarrow 3'$ direction. The value of $k_\alpha$ is 5 kcal mol$^{-1}$ rad$^{-2}$ for angles involving base beads and 20 kcal mol$^{-1}$ rad$^{-2}$ for all other angles.

The potential to model the hydrogen-bonding network of RNA is given by

$$U_{\text{HB}} = U_{\text{HB}}^0 \exp(-u), \tag{S4}$$

where

$$u = 5.0(r-r_0)^2+1.5\big\{(\theta_1-\theta_{1,0})^2+(\theta_2-\theta_{2,0})^2\big\}+0.15\big\{(\psi_0-\psi_{0,0})^2+(\psi_1-\psi_{1,0})^2+(\psi_2-\psi_{2,0})^2\big\}.$$

The value of $U^0_{\mathrm{HB}}$ is -2.9 kcal·mol$^{-1}$. The parameters $r_0$, $\theta_{1,0}$, $\theta_{2,0}$, $\psi_{0,0}$, $\psi_{1,0}$, and $\psi_{2,0}$ for all the hydrogen bonds are calculated from the coarse-grained model of the two crystal structures of SAM-III riboswitch (PDB ID: 3E5C, 6C27).[1–4] The canonical hydrogen bonds included in the force field are shown in the secondary structures (see Fig.1b,c in the Main text). The additional hydrogen bonds and tertiary stacking interactions are listed in Table S2,S3. The definitions of $r$, $\theta_1$, $\theta_2$, $\psi_0$, $\psi_1$, and $\psi_2$ are provided in the work of Denesyuk et al.[2] The hydrogen-bond network of the system is identified using the two crystal structures and the RNApdbee 2.0 server.[5]

The sequence-dependent stacking interactions between two consecutive nucleotides ($U_{\mathrm{ST}}$) are given by:

$$U_{\mathrm{ST}} = \frac{U^0_{\mathrm{ST}}}{1 + k_r(r-r_0)^2 + k_\phi(\phi_1-\phi_{1,0})^2 + k_r(\phi_2-\phi_{2,0})^2} \tag{S5}$$

where, $k_r$ is 1.4 Å$^{-2}$ and $k_\phi$ is 4.0 rad$^{-2}$. The equilibrium parameters $r_0$, $\phi_{1,0}$, and $\phi_{2,0}$ are taken from the coarse-grained structure of an ideal A-form RNA helix. The values of $U^0_{\mathrm{ST}}$ and all the definitions of $r$, $\phi_1$, and $\phi_2$ are defined in the original work of Denesyuk et al.[1]

The tertiary stacking interaction between two non-adjacent nucleotide bases ($U_{\mathrm{TST}}$) is given by

$$U_{\mathrm{TST}} = \frac{U^0_{\mathrm{TST}}}{1+u} \tag{S6}$$

where,

$$u = 5.0(r-r_0)^2+1.5\big\{(\theta_1-\theta_{1,0})^2+(\theta_2-\theta_{2,0})^2\big\}+0.15\big\{(\psi_0-\psi_{0,0})^2+(\psi_1-\psi_{1,0})^2+(\psi_2-\psi_{2,0})^2\big\}.$$

The value of $U^0_{\mathrm{TST}}$ is taken to be -4.75 kcal·mol$^{-1}$. The definitions of the other variables in the potential are the same as the hydrogen bonding interaction potential used in the work of Denesyuk et al.[2]

The electrostatic interaction between any pair of particles is computed using the Coulomb potential given by:

$$U_{\mathrm{EL}} = \frac{Z_i Z_j}{4\pi\epsilon_0\epsilon_r r_{ij}} \tag{S7}$$

where $Z_i$ and $Z_j$ are the charges on the particles $i$ and $j$, $\epsilon_0$ is the vacuum permittivity, and $\epsilon_r(T)$ is the dielectric constant of water, which is a function of temperature, and is given by

$$\epsilon_r(T) = 87.740 - 0.4008T + 9.398 \times 10^{-4}T^2 + 1.410 \times 10^{-6}T^3 \tag{S8}$$

The excluded volume interactions between the beads are modeled using a modified LJ potential given by

$$U_{\mathrm{EV}} = \begin{cases} \epsilon_{ij}\left[\left(\dfrac{d_s}{r + d_s - \rho_{ij}}\right)^{12} - 2\left(\dfrac{d_s}{r + d_s - \rho_{ij}}\right)^{6} + 1\right] & \text{if } r \leq d_{ij} \\ 0 & \text{if } r > 0 \end{cases} \tag{S9}$$

where $d_s$ is the diameter of the smallest ion present in the system, $\rho_{ij} = R_i + R_j$, $\epsilon_{ij} = \sqrt{\epsilon_i\epsilon_j}$. The values of $R_i$ and $\epsilon_i$ for phosphate (P), sugar (S), and bases (A, G, C, and U) are taken from the work of Denesyuk et al.[2] The values of $R_i$ and $\epsilon_i$ for the ions are given in Table S4.

#### Simulation Protocol

We used the three interaction site (TIS-RNA)[1,2] model to investigate the *apo*↔R conformational transition in SAM-III riboswitch. The initial structure for the simulations was prepared by combining the sequence information from both crystal structures to generate a common 60-nucleotide sequence. We performed Langevin dynamics simulations using OpenMM[6] at different temperatures ($T$ = 295 K, 310 K, 338 K, and 368 K) using a cubic box of length 200 Å. The water solvent is implicitly modeled. Simulations were performed at various ion concentrations, $[\mathrm{Mg}^{2+}] = 2, 5, 8$ mM with a fixed background $[\mathrm{K}^{+}] = 150$ mM. The $K^+$, $Mg^{2+}$, and $Cl^-$ ions were randomly inserted into the box to obtain the desired physiological ion concentrations while maintaining charge neutrality. Simulation duration for different systems is listed in S1. We discarded the initial 1 $\mu$s of the data from each CG trajectory for analysis. The equation of motion for the particles in the simulation is given by

$$m_i \ddot{\boldsymbol{r}}_{\boldsymbol{i}} = -m_i \gamma \dot{\boldsymbol{r}}_{\boldsymbol{i}} + \boldsymbol{F}_i + \boldsymbol{\Gamma}_i \tag{S10}$$

where, $\boldsymbol{r_i}$ and $\gamma$ are coordinates and friction coefficient of the $i$th particle, respectively. The deterministic force on the particle $i$ is given by $\boldsymbol{F}_i = -\dfrac{\partial U_{\mathrm{TIS}}(\boldsymbol{r_i})}{\partial \boldsymbol{r_i}}$, and $\boldsymbol{\Gamma}_i$ is an uncorrelated random force with a white noise spectrum. The autocorrelation function of the random force in discretized form is $\big\langle \boldsymbol{\Gamma}(t) \cdot \boldsymbol{\Gamma}(t + nh) \big\rangle = \frac{2\gamma m_i k_{\mathrm{B}} T}{h} \delta_{0,n}$, where $n = 0, 1, ...$ and $\delta_{0,n}$ is the Kronecker delta function. We have used the *LangevinIntegrator* module in OpenMM to integrate the equations of motion. For better conformational sampling, we used a low friction coefficient ($\gamma = 0.01$ ps$^{-1}$)[7] with an integration time step of 5 fs. The Particle Mesh Ewald (PME) algorithm was used to compute the long-range Coulomb interactions. We used the TIS2AA program[8] to generate all-atom representations of the coarse-grained structures. We used VMD[9] for visualization and representation of the conformations.

### Data Analysis

#### Calculation of the Free Energy Difference Between A2 and R

ITC experiments combined with kinetic modeling have reported the conformational equilibrium constant, $K_{\mathrm{ISO}}$, between the $\mathrm{ISO}_{\mathrm{SMK}}$ and $\mathrm{PRIMED}_{\mathrm{SMK}}$ states of the SAM-responsive $S_{\mathrm{MK}}$ riboswitch. The detailed calculation of $K_{\mathrm{ISO}}$ is provided in the work of Wilson et al.[10] In our simulations, $\mathrm{ISO}_{\mathrm{SMK}}$ and $\mathrm{PRIMED}_{\mathrm{SMK}}$ correspond structurally to the A2 and R states, respectively. Thus, we used the reported value $K_{\mathrm{ISO}} \approx 5.3$ to estimate the experimental free-energy difference between A2 and R.

Because $K_{\mathrm{ISO}}$ is defined for the conformational equilibrium between $\mathrm{PRIMED}_{\mathrm{SMK}} \rightleftharpoons \mathrm{ISO}_{\mathrm{SMK}}$ (which is $\mathrm{R} \rightleftharpoons \mathrm{A2}$, in simulations), it corresponds in our notation to $K_{\mathrm{ISO}} = P_{\mathrm{A2}}/P_{\mathrm{R}}$. Therefore, the free energy of R relative to A2 was calculated as

$$\Delta G^{\mathrm{exp}}_{\mathrm{A2} \rightleftharpoons \mathrm{R}} = G_{\mathrm{R}} - G_{\mathrm{A2}} = k_{\mathrm{B}}T \ln K_{\mathrm{ISO}}. \tag{S11}$$

Using $K_{\mathrm{ISO}} = 5.3$, we obtained $\Delta G^{\mathrm{exp}}_{\mathrm{A2} \rightleftharpoons \mathrm{R}} \approx 1.67\ k_{\mathrm{B}}T$. At the experimental temperature, $T_{\mathrm{exp}} = 288$ K, this value corresponds to $\Delta G^{\mathrm{exp}}_{\mathrm{A2} \rightleftharpoons \mathrm{R}} \approx 0.96$ kcal mol$^{-1}$.

+

### Computation of Transition Path Probabilities

We observed four distinct transition pathways, and $n_i$ denotes the number of transitions observed along the pathway $i$. The probability of each pathway, $P_i$, at a given [$\mathrm{Mg}^{2+}$] was calculated by normalizing its transition count by the total number of transitions across all four pathways and is given by

$$P_i = \frac{n_i}{\sum_{j=1}^{4} n_j}, \qquad i = 1, \ldots, 4, \tag{S12}$$

such that $\sum_{i=1}^{4} P_i = 1$.

### Computation of Contact Maps

To compute the average contact map, we determined the pairwise distances between all the RNA CG beads, excluding the bead pairs that are separated by less than three nucleotides apart from each other in the RNA sequence. A pair of beads was considered to be in contact if the distance between them was less than the cutoff distance, $r_{\mathrm{cut}} = 15$ Å.[2,11] The contact map was then averaged over all conformations in the corresponding sub-ensemble.

#### Computation of Spatial Distribution of Ions Around RNA

We used the volmap plugin implemented in the VMD software[9] to generate spatial density maps that illustrate the distribution of $Mg^{2+}$ ions around RNA nucleotides. Before calculating the density map, we aligned the RNA beads from all conformations to the final conformation of the CG trajectory. The coordinates of the $Mg^{2+}$ ions in each simulation frame were adjusted accordingly. The alignment process utilized the Kabsch method,[12] which is implemented in VMD. In the density calculations, $Mg^{2+}$ ions were modeled as spheres with their van der Waals radii. A grid with a size of 0.01 $nm^3$ was constructed to partition the space surrounding the RNA nucleotides. A grid point was considered as occupied and assigned a value of 1 if the sphere corresponding to a $Mg^{2+}$ ion overlapped with it; otherwise, it was assigned a value of 0. The occupation values at each point on the grid were averaged across all frames of the trajectory to obtain spatial density map. VMD was also utilized for structural superposition[13] and for image rendering.[14]

Table S1: Description of CG simulation systems. All the system contains 723 $K^+$ ions corresponding to $[K^+]$ = 150 mM.

| **System** | $T_{\mathbf{sim}}$ **(K)** | $\mathbf{[Mg^{2+}]}$ **concentration (mM)** | **Number of ions** ($Mg^{2+}$, $Cl^-$) | **Simulation duration** ($\mu$s) **per replicate (No. of replicates)** | *apo*↔*holo* **transition count** |
|---|---|---|---|---|---|
| SAM-III | 295 | 5 | 24, 711 | ≈ 20 (18) | 15 |
| SAM-III | 295 | 8 | 39, 741 | ≈ 20 (8) | 7 |
| SAM-III | 310 | 2 | 10, 683 | ≈ 30 (32) | 84 |
| SAM-III | 310 | 5 | 24, 711 | ≈ 30 (20) | 65 |
| SAM-III | 310 | 8 | 39, 741 | ≈ 30 (16) | 68 |
| SAM-III | 338 | 5 | 24, 711 | ≈ 10 (8) | 39 |
| SAM-III | 338 | 8 | 39, 741 | ≈ 10 (8) | 34 |
| SAM-III | 368 | 5 | 24, 711 | ≈ 10 (2) | NA |
| Mutated SAM-III | 295 | 5 | 24, 711 | ≈ 10 (4) | NA |

Table S2: Non-canonical H-bonds in base-base, sugar-sugar, phosphate-sugar and base-base tertiary stacks present in the *apo* (PDB ID: 6C27) crystal structure.[4] Canonical base-base H-bonds are shown in the secondary structure (Main text Fig.1b).

| Base-Base Non-canonical | Sugar-Sugar | Phosphate-Sugar | Base-Base Tertiary Stack |
|---|---|---|---|
| U9-C39(1)<br>C13-A17(1)<br>G35-G38(2) | U9-G35 | U10-A34 | U9-G35<br>A6-G43 |

Table S3: Non-canonical H-bonds in base-base, base-sugar, phosphate-sugar, and base-base tertiary stacks present in the *holo* (PDB ID: 3E5C) crystal structure.[3] Canonical base-base H-bonds are shown in the secondary structure (Main text Fig.1c).

| Base-Base Non-canonical | Base-Sugar | Phosphate-Sugar | Base-Base Tertiary Stack |
|---|---|---|---|
| A15-G43(2)<br>G38-G43(2)<br>A45-G55(1) | U44-G14<br>A46-G55 | U44-A15 | A36-G38<br>G14-A36 |

Table S4: RNA[2] and ions[15,16] parameters used in the CG simulations.

| Type | $R_i$ (Å) | $m_i$ (amu) | $\epsilon_i$ (kcal mol$^{-1}$) | $Z_i$ |
|---|---|---|---|---|
| P | 2.1 | 62.971 | 0.2 | -1 |
| S | 2.9 | 131.108 | 0.2 | 0 |
| A | 2.8 | 134.119 | 0.2 | 0 |
| G | 3.0 | 150.118 | 0.2 | 0 |
| C | 2.7 | 110.094 | 0.2 | 0 |
| U | 2.7 | 111.079 | 0.2 | 0 |
| $Mg^{2+}$ | 1.353 | 24.305 | 0.009 | +2 |
| $K^+$ | 1.590 | 39.098 | 0.279 | +1 |
| $Cl^-$ | 2.760 | 35.453 | 0.012 | -1 |

Table S5: Description of nucleotides of sub-parts of SAM-III

| **Region** | **Nucleotides** |
| --- | --- |
| *apo* (specific) | G1-C47 |
| *holo* (specific) | G8-U60 |
| P0 | G1-G8, C40-C47 |
| P1 | G8-C13, G55-U60 |
| P2 | A16-G19, C39-U42 |
| P3 | U21-A34 |
| P4 | C47-G54 |
| P5 | U10-A20 |
| $3WJ^{apo}$ | A7-C11, G20-G23, C33-U41 |
| $3WJ^{holo}$ | C13-A16, U43-C47, G54-G55 |

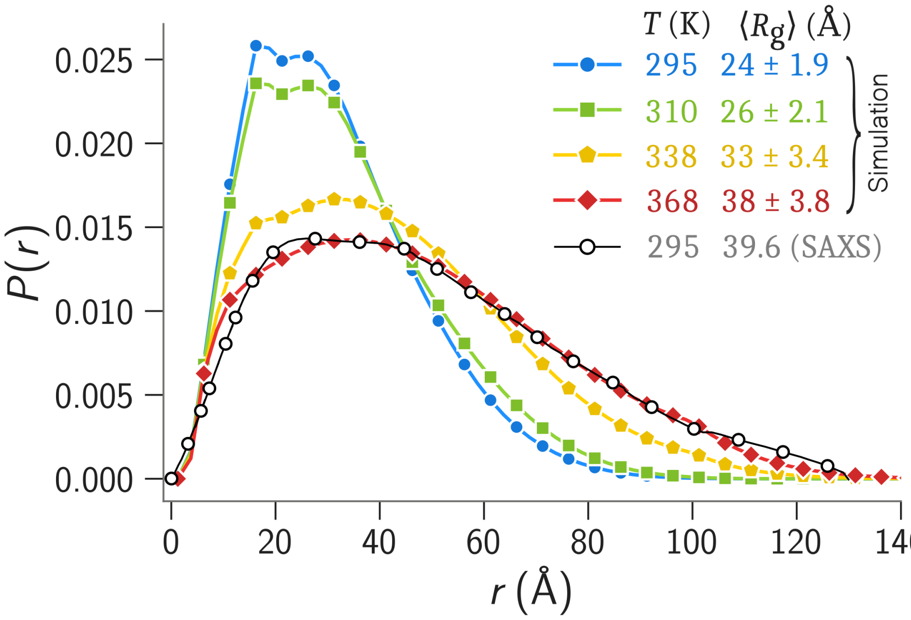


Figure S1: Pair distance distribution (PDD) function of SAM-III(59) from SAXS experiments (black empty circles) and simulation trajectories at 5 mM [$Mg^{2+}$] and $T_{\rm sim}$ = 295 K (blue solid circles), $T_{\rm sim}$ = 310 K (green solid squares), $T_{\rm sim}$ = 338 K (golden solid pentagons) and $T_{\rm sim}$ = 368 K (red solid diamonds).

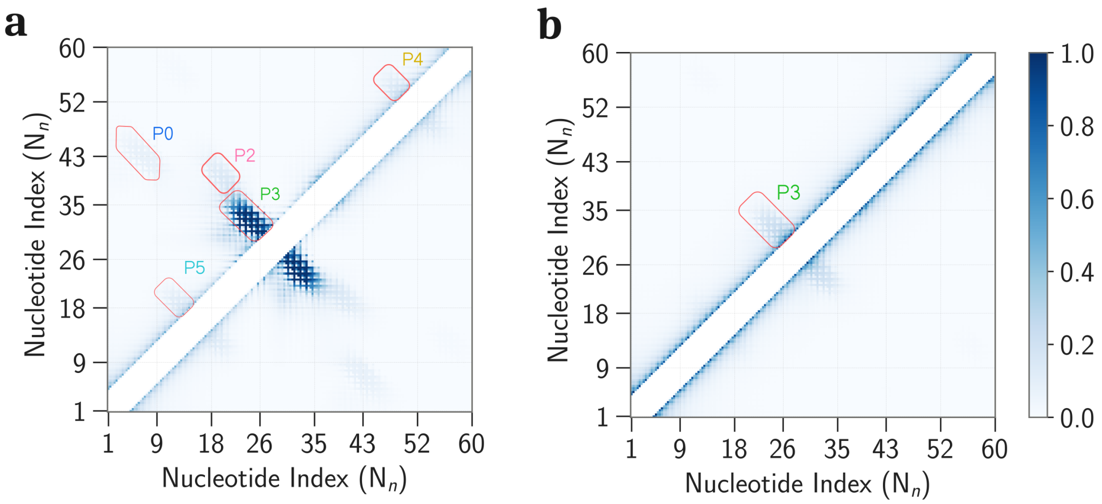


Figure S2: The average contact map of SAM-III conformational ensemble from simulations at [$Mg^{2+}$] = 5 mM and (a) $T_{\rm sim}$ = 338 K, and (b) 368 K. At 338 K, P3 helix is most stable, while P0, P2, P4, P5 helices are marginally stable. At 368 K, only P3 helix is marginally stable.

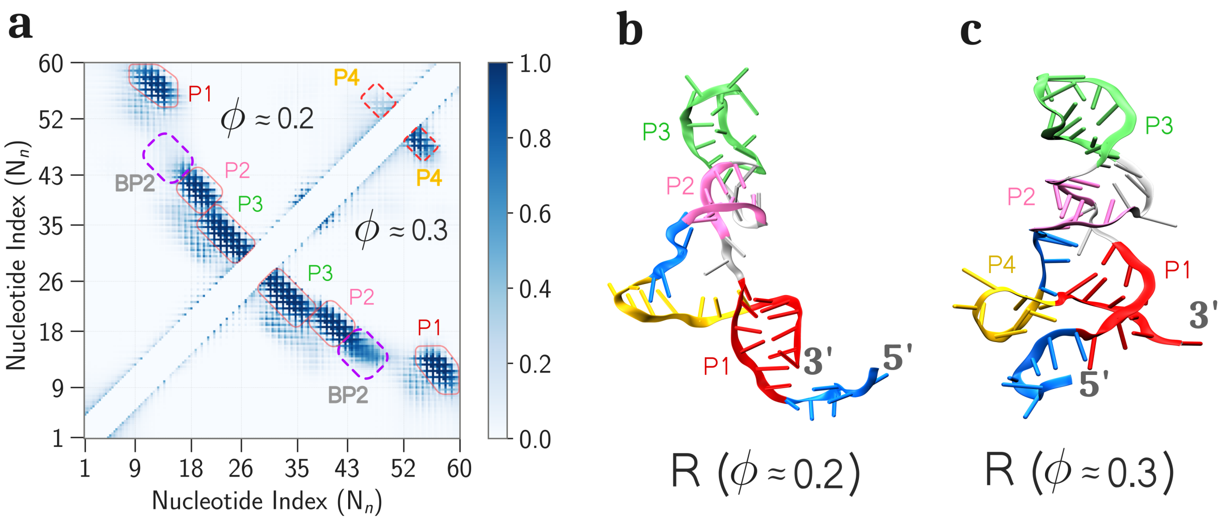


Figure S3: **(a)** The average contact map of R($\phi \approx 0.2$) (upper diagonal) and R ($\phi \approx 0.3$) (lower diagonal) states. Cartoon representation of tertiary structure of **(b)** R ($\phi \approx 0.2$) and **(c)** R ($\phi \approx 0.3$) states.

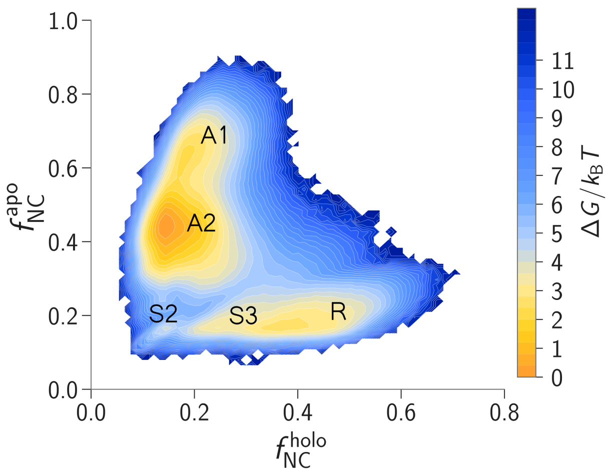


Figure S4: 2D FES projected on the fraction of native contacts ($f_{\mathrm{NC}}^{\mathrm{holo}}$) and ($f_{\mathrm{NC}}^{\mathrm{apo}}$) at $T_{\mathrm{sim}} =$ 310 K, [$K^+$] = 150 mM, and [$Mg^{2+}$] = 2 mM.

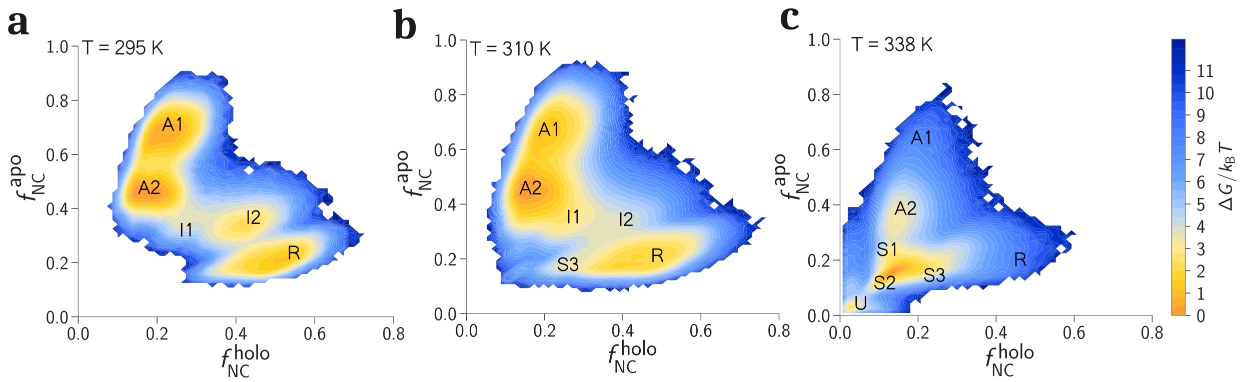


Figure S5: 2D FES projected on the fraction of native contacts $f_{\rm NC}^{\rm holo}$ and $f_{\rm NC}^{\rm apo}$ at $[K^+] = 150$ mM and $[Mg^{2+}] = 5$ mM. FES at **(a)** $T_{\rm sim} = 295$ K **(b)** $T_{\rm sim} = 310$ K and **(c)** $T_{\rm sim} = 338$ K highlights the shift of population towards semi-folded states with increasing temperature. U refers to an unfolded ensemble where even P3 is not properly formed.

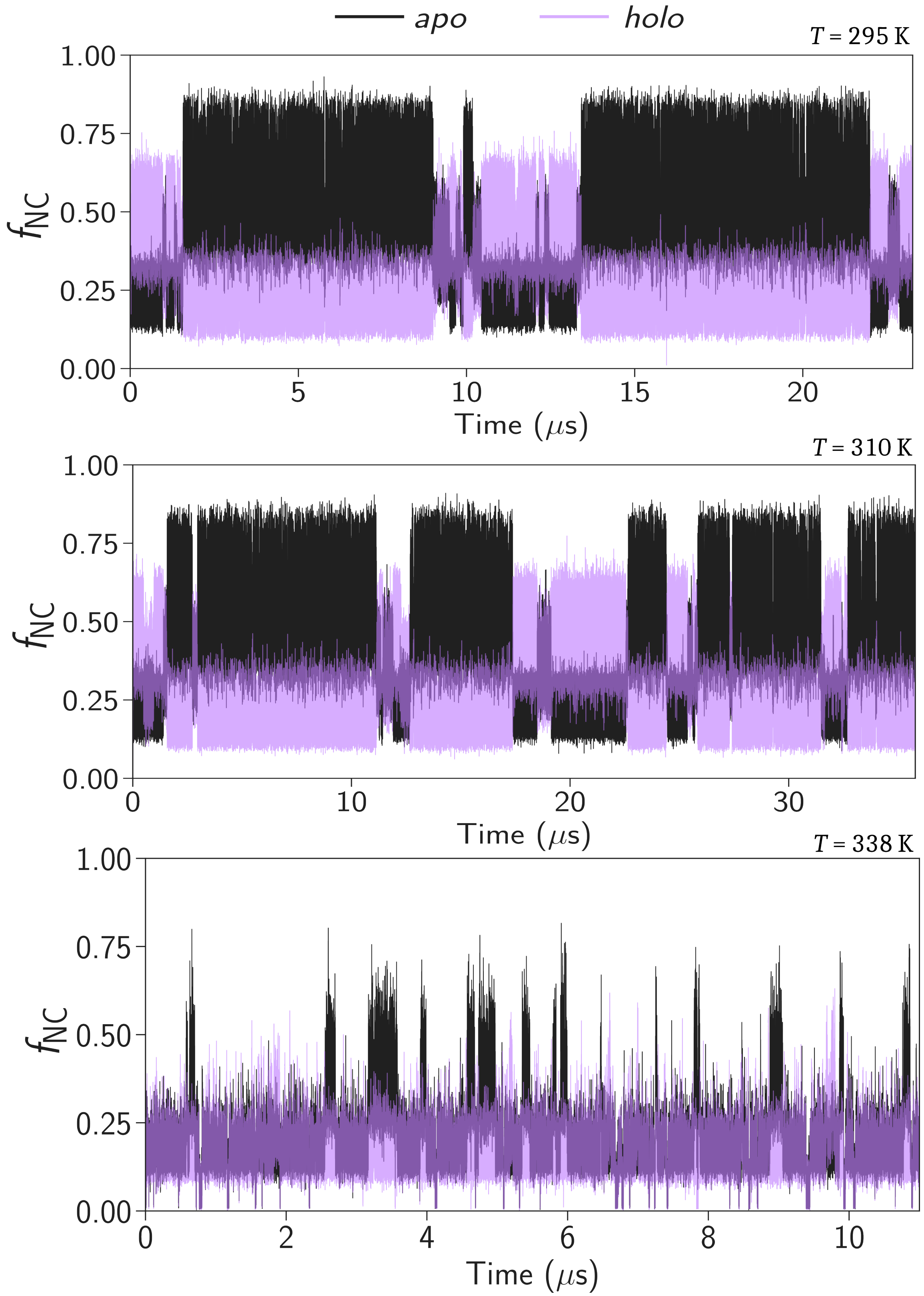


Figure S6: Representative trajectory showing multiple transitions between the A1 and R states at temperatures $T_{\mathrm{sim}} = 295$K, 310K, and 338K, respectively, and $[\mathrm{Mg}^{2+}] = 5$ mM.

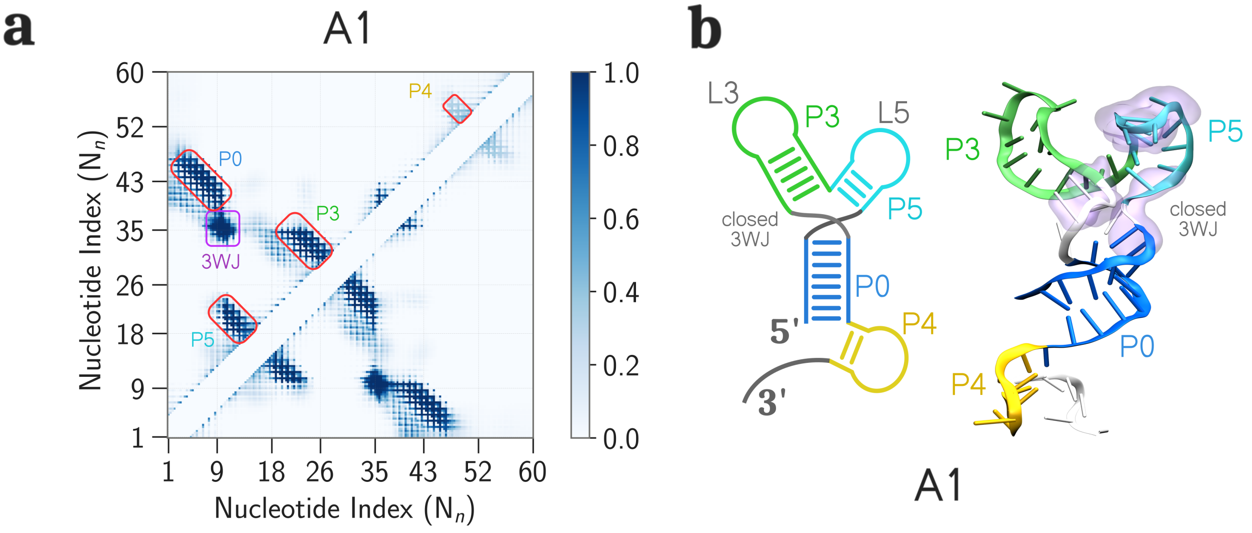


Figure S7: **(a)** The average contact map of A1 state. Both the upper and lower diagonals of the contact map are from simulations. **(b)** Schematic and cartoon representation of tertiary structure of A1 state. Nucleotides involved in the native contacts formation associated with 3WJ in the *apo* crystal structure[4] are highlighted in transparent surfaces (pale violet).

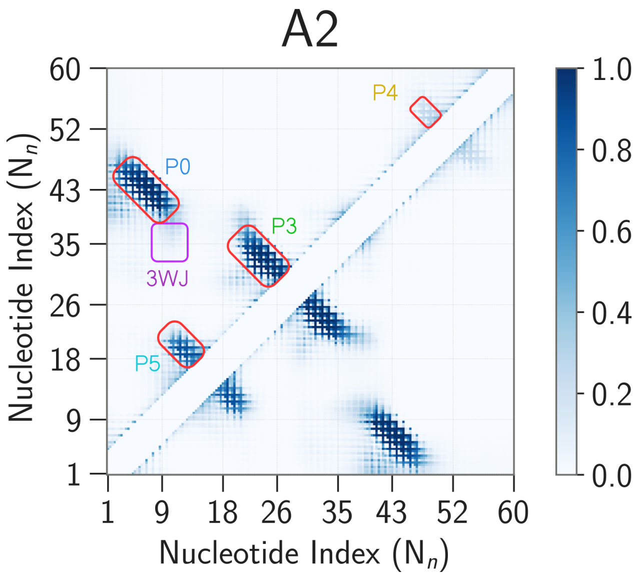


Figure S8: The average contact map of A2 state. The native contacts associated with the 3WJ are missing. Both the upper and lower diagonals of the contact map are from simulations.

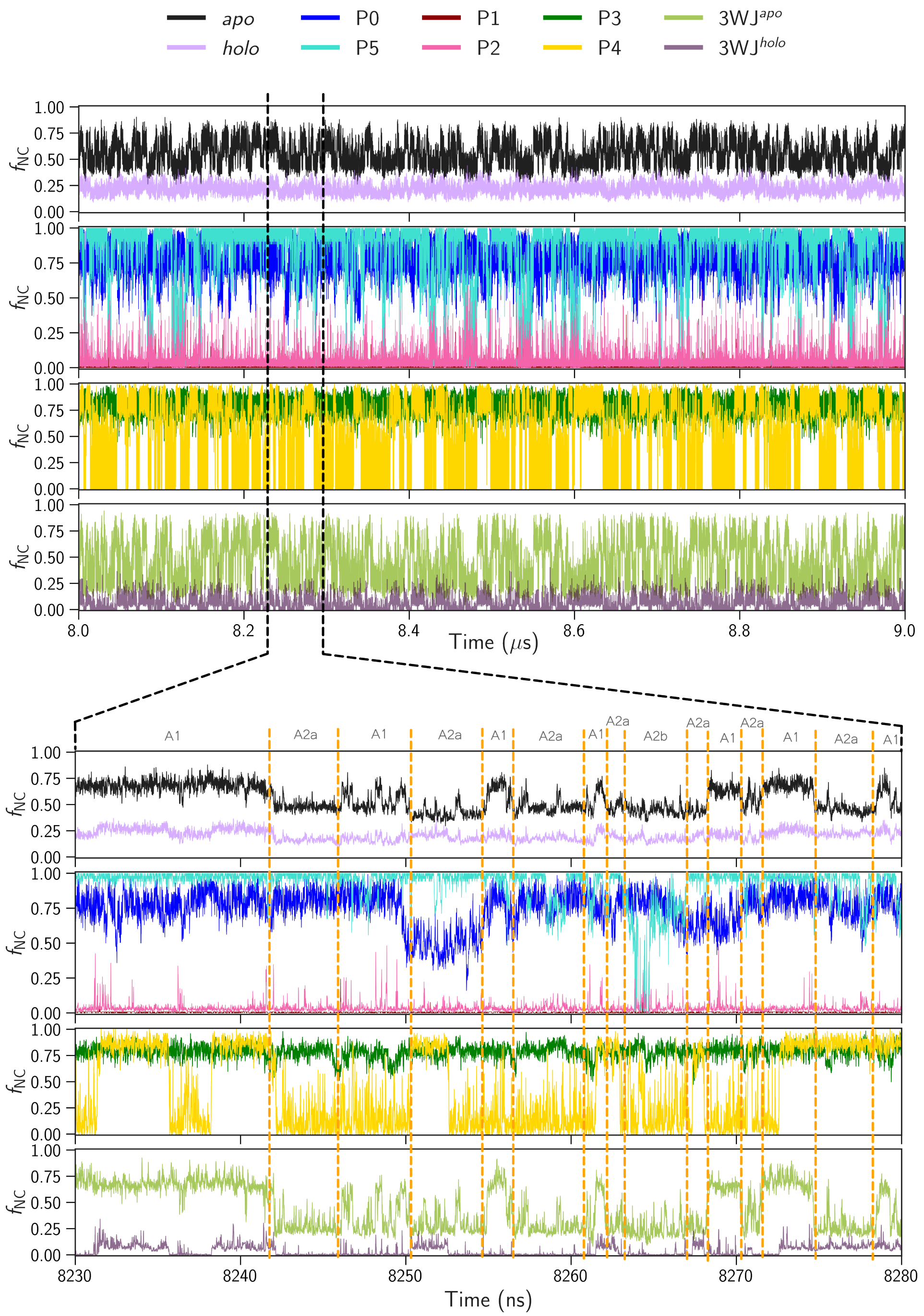


Figure S9: Representative trajectory segment showing multiple transitions between the A1 and A2 states highlighting the breathing motion of the $3WJ^{apo}$.

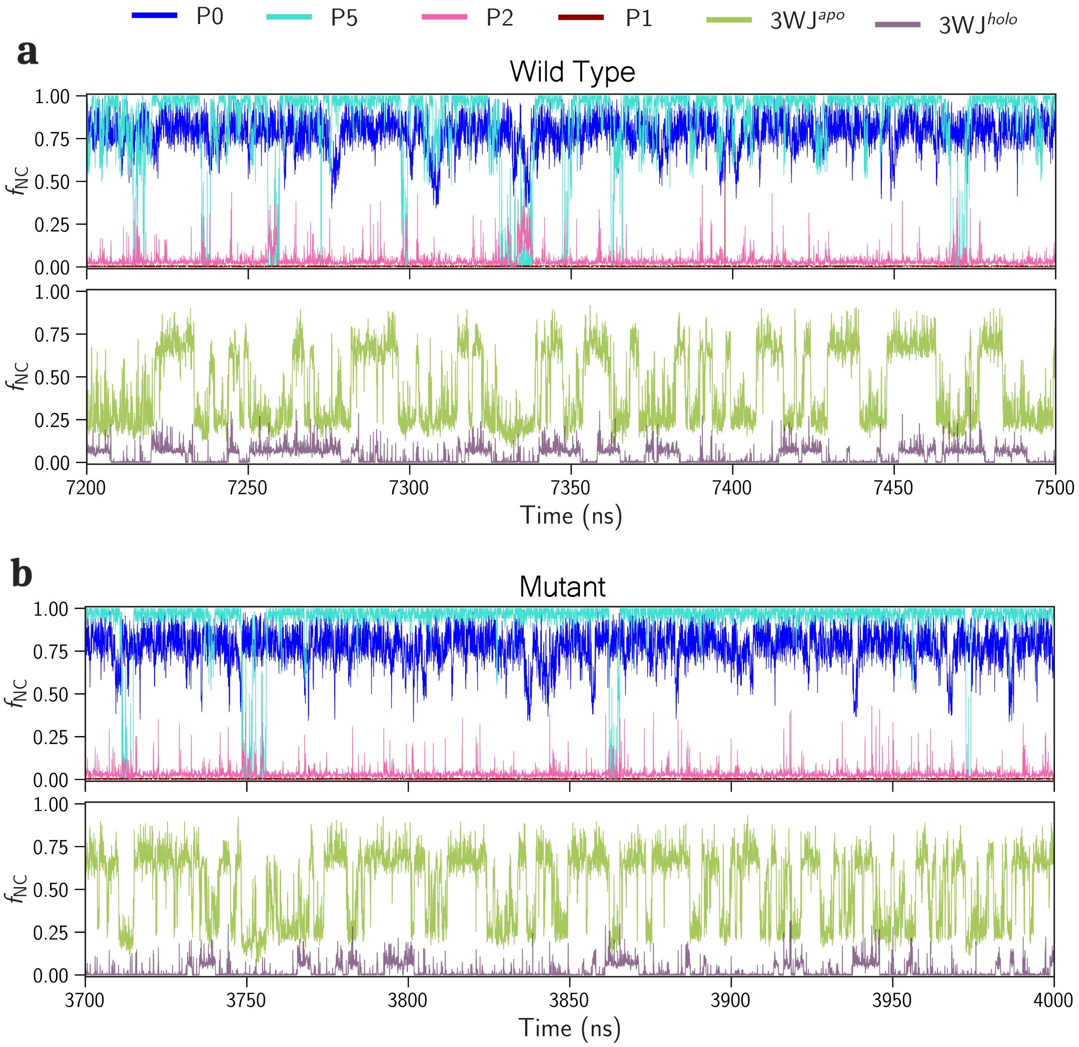


Figure S10: Representative trajectory segment for **(a)** Wild type and **(b)** Mutated structure (U10C and A20G) at $T_{\text{sim}} = 295$ K, $[\text{K}^+] = 150$ mM and $[\text{Mg}^{2+}] = 5$ mM. The P5 helix and 3WJ of *apo* structure shows less disruption in the mutated structure.

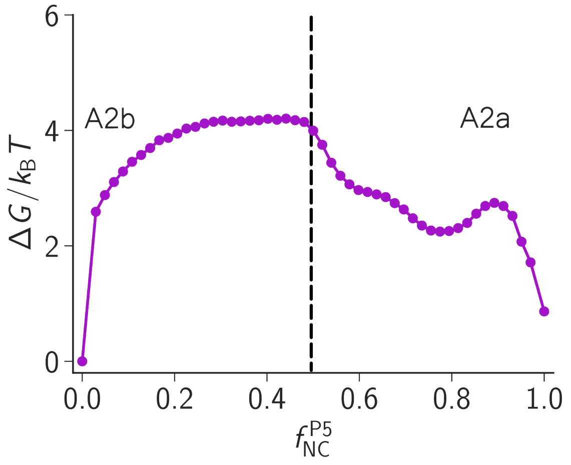


Figure S11: FES projected on fraction of native contacts of P5 ($f_{\text{NC}}^{\text{P5}}$). The FES is computed using the A2 conformational ensemble, showing the existence of A2a and A2b microstates.

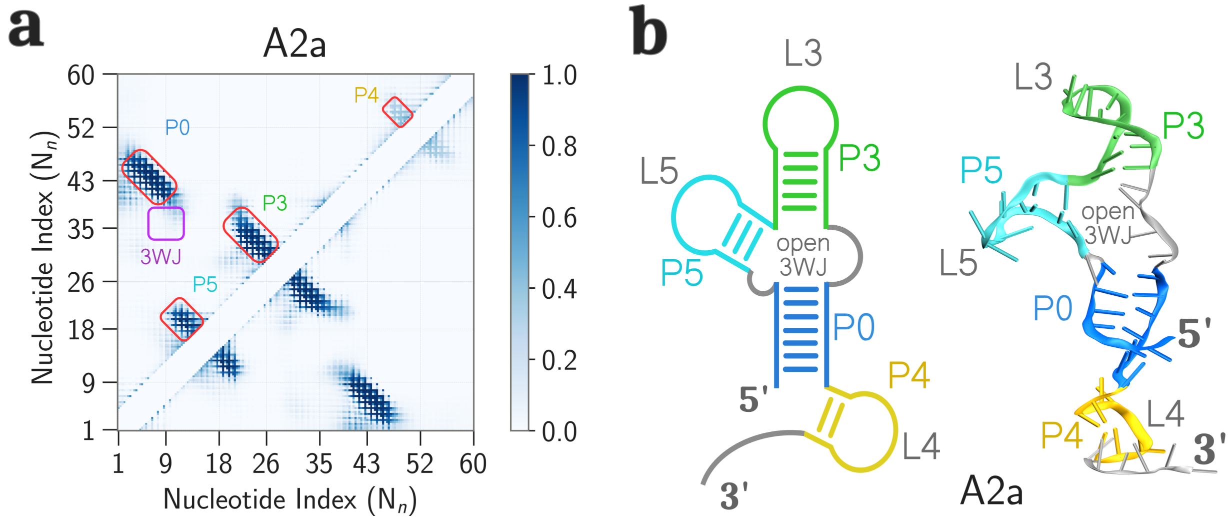


Figure S12: **(a)** The average contact map of A2a state. Both the upper and lower diagonals of the contact map are from simulations. **(b)** Schematic and cartoon representation of tertiary structure of A2a state.

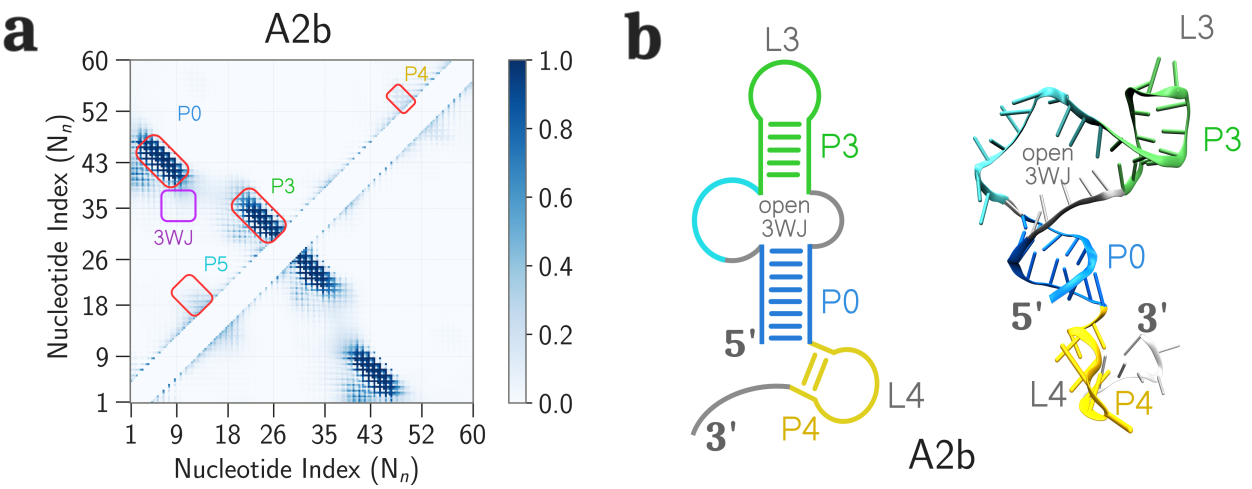


Figure S13: The average contact map of A2b state. Both the upper and lower diagonals of the contact map are from simulations. **(b)** Schematic and cartoon representation of tertiary structure of A2b state.

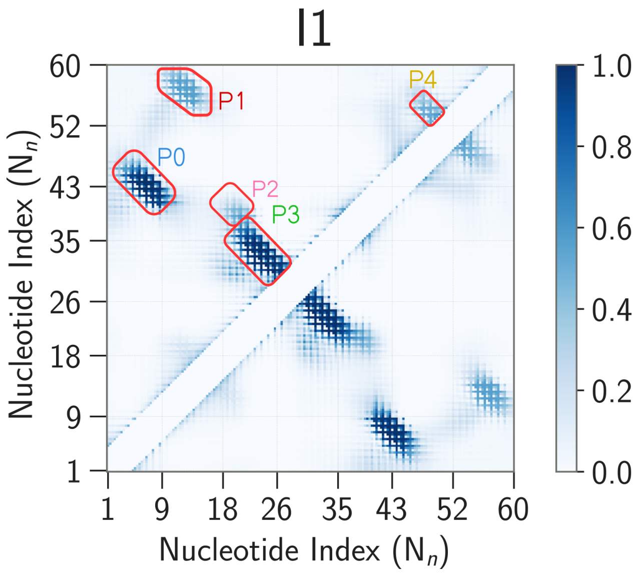


Figure S14: The average contact map of I1 state. Both the upper and lower diagonals of the contact map are from simulations.

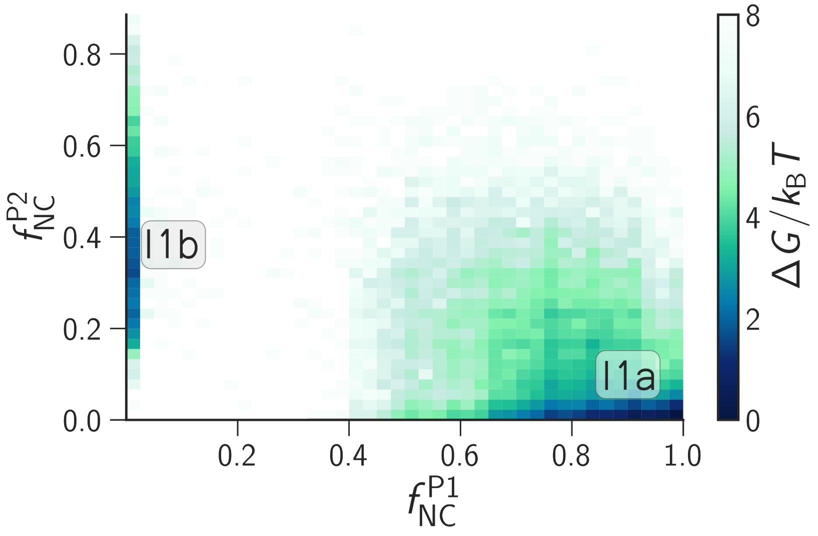


Figure S15: 2D FES projected on fraction of native contacts of P1 ($f_{\mathrm{NC}}^{\mathrm{P1}}$) and P2 ($f_{\mathrm{NC}}^{\mathrm{P2}}$) using the I1 conformational ensemble showing the existence of I1a and I1b microstates.

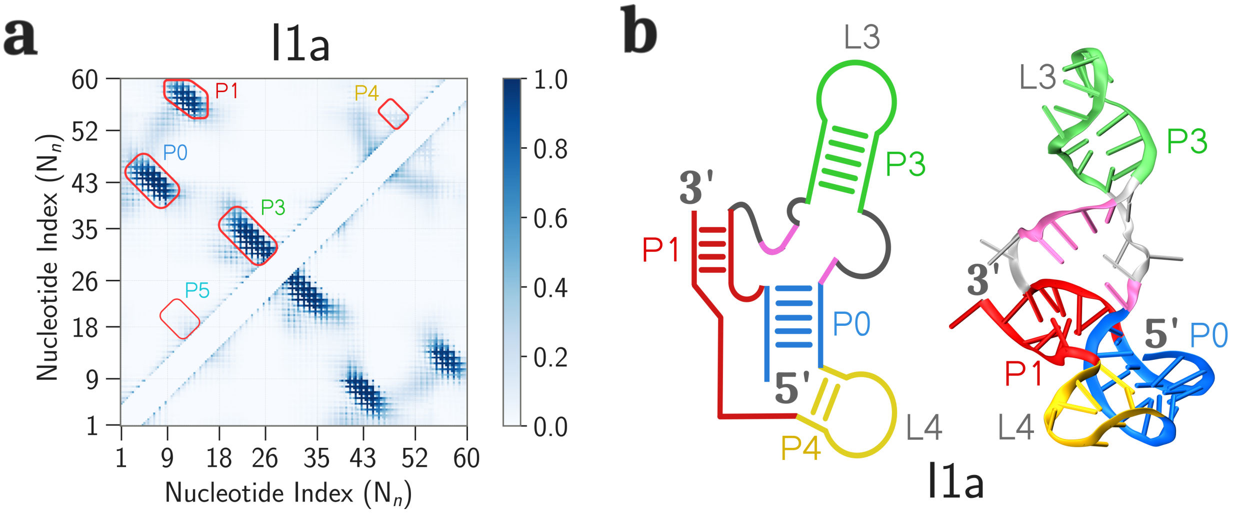


Figure S16: **(a)** The average contact map of I1a state. Both the upper and lower diagonals of the contact map are from simulations. **(b)** Schematic and cartoon representation of tertiary structure of I1a state.

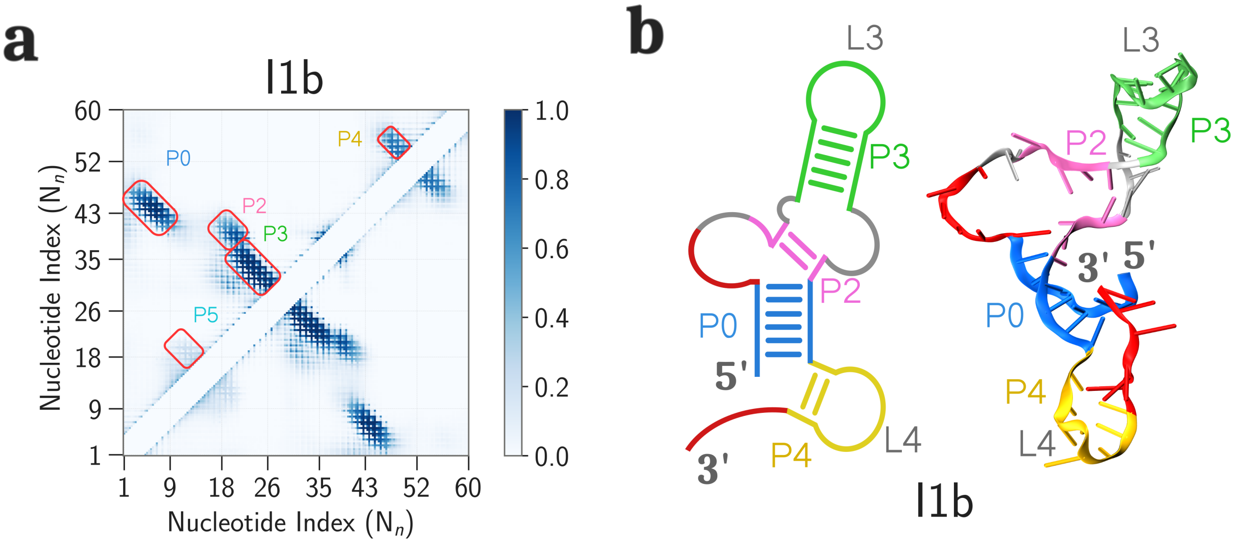


Figure S17: **(a)** The average contact map of I1b state. Both the upper and lower diagonals of the contact map are from simulations. **(b)** Schematic and cartoon representation of tertiary structure of I1b state.

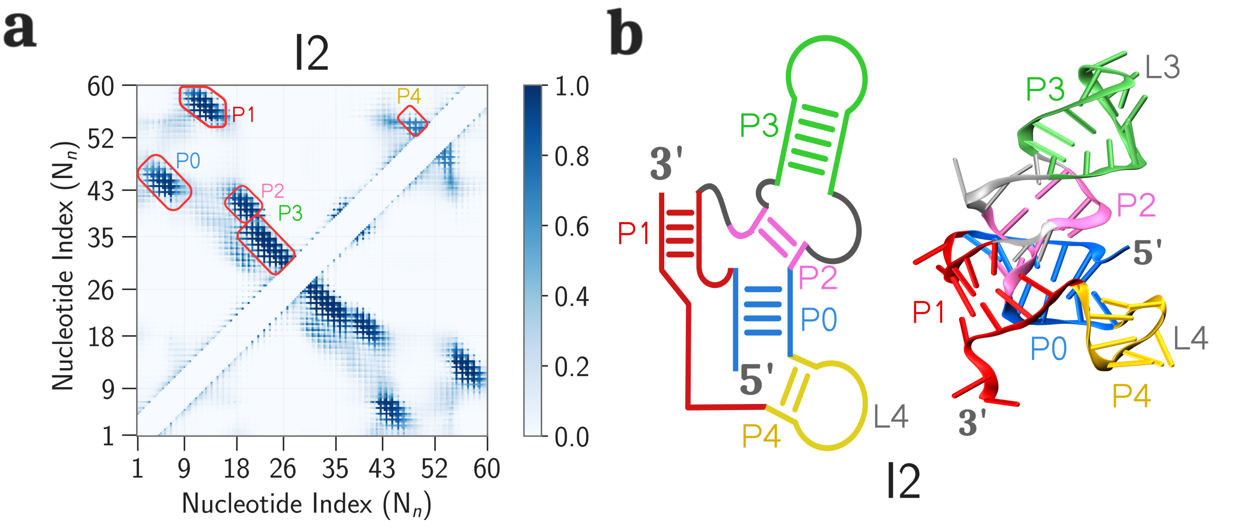


Figure S18: **(a)** The average contact map of I2 state. Both the upper and lower diagonals of the contact map are from simulations. **(b)** Schematic and cartoon representation of tertiary structure of I2 state.

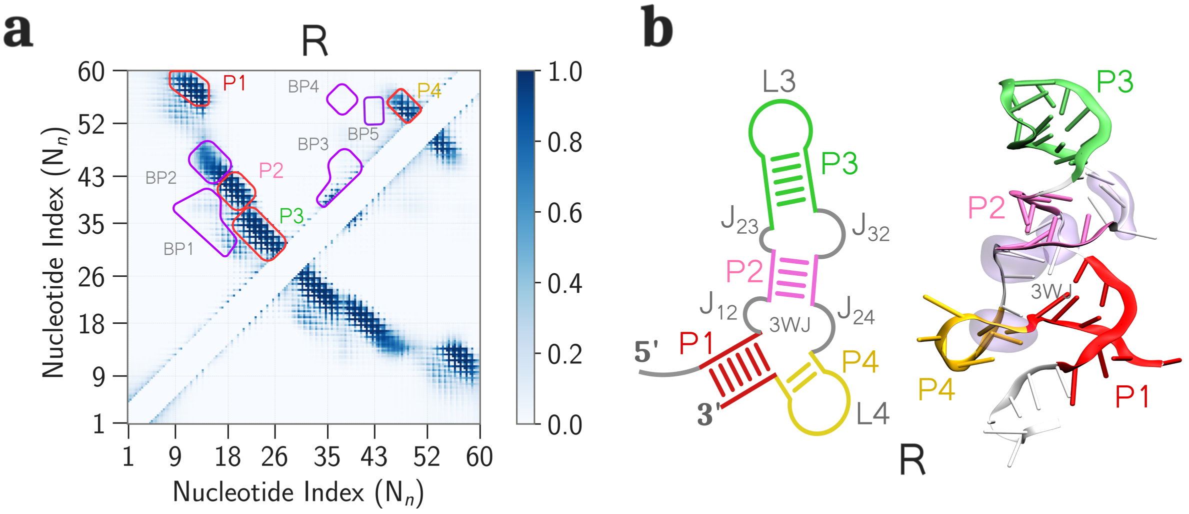


Figure S19: **(a)** The average contact map of R state. Both the upper and lower diagonals of the contact map are from simulations. **(b)** Schematic and cartoon representation of the tertiary structure of the R state. Nucleotides involved in the native contacts formation associated with binding pocket (BP1–BP5) in the *holo* crystal structure[3] are highlighted in transparent surfaces (pale violet).

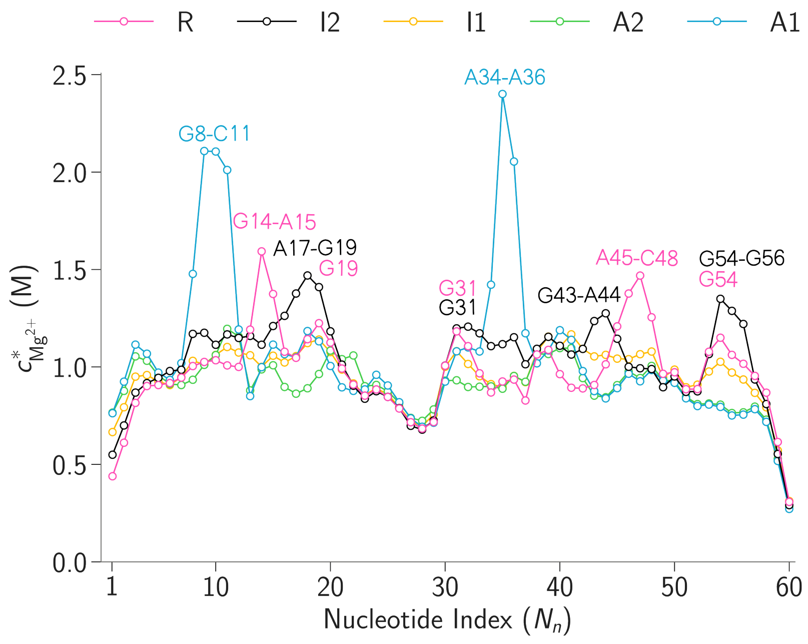


Figure S20: The local ion ($c^*_{Mg^{2+}}$) profile for the five states around phosphate oxygen atoms (OP) of all the nucleotides at $[Mg^{2+}] = 8$ mM and $T_{sim} = 310$ K.

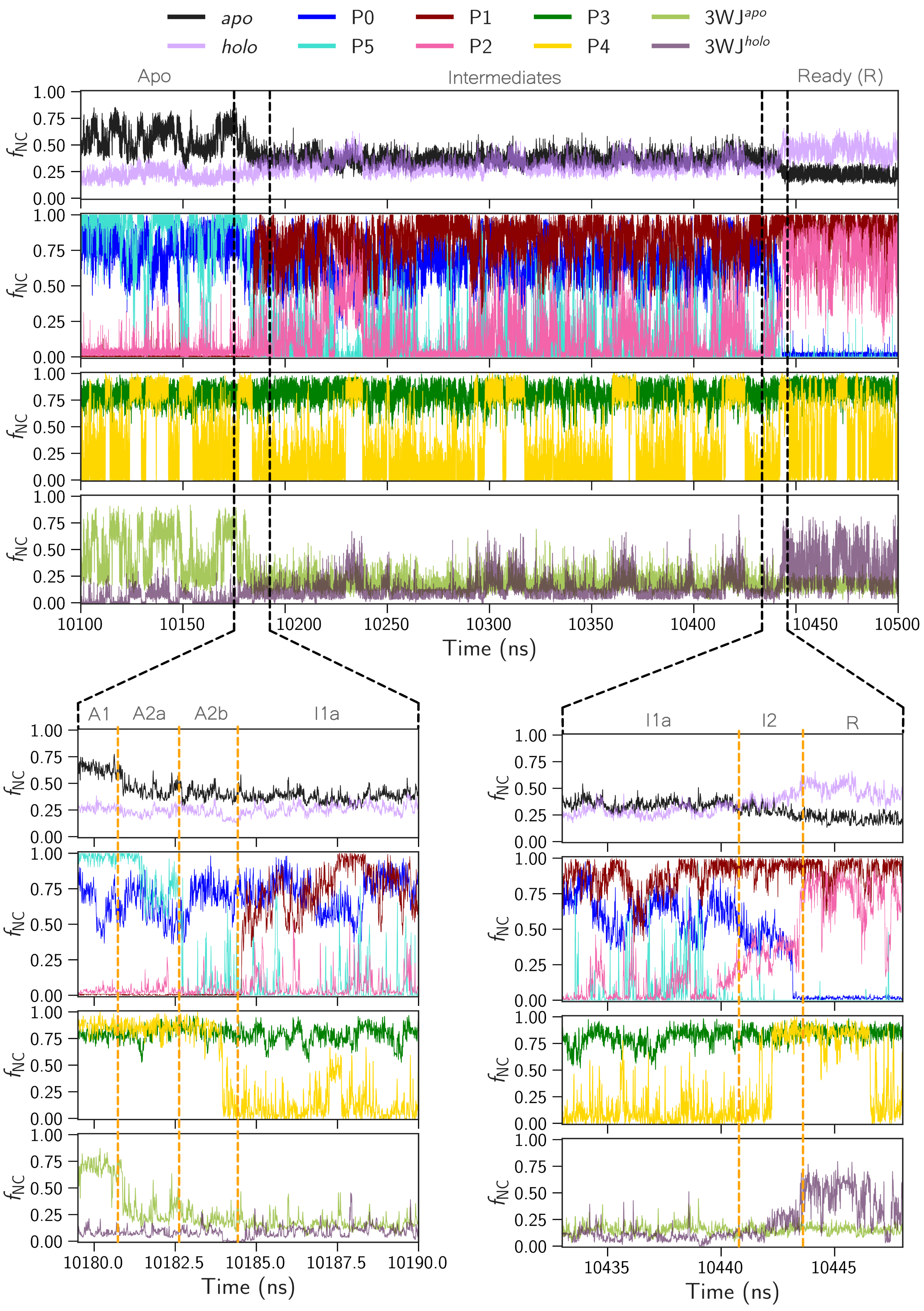


Figure S21: Representative trajectory segment for transition pathway of A1 → A2a → A2b → I1a → I2 → R. The different $f_{NC}$ components are shown in four rows for transition identification.

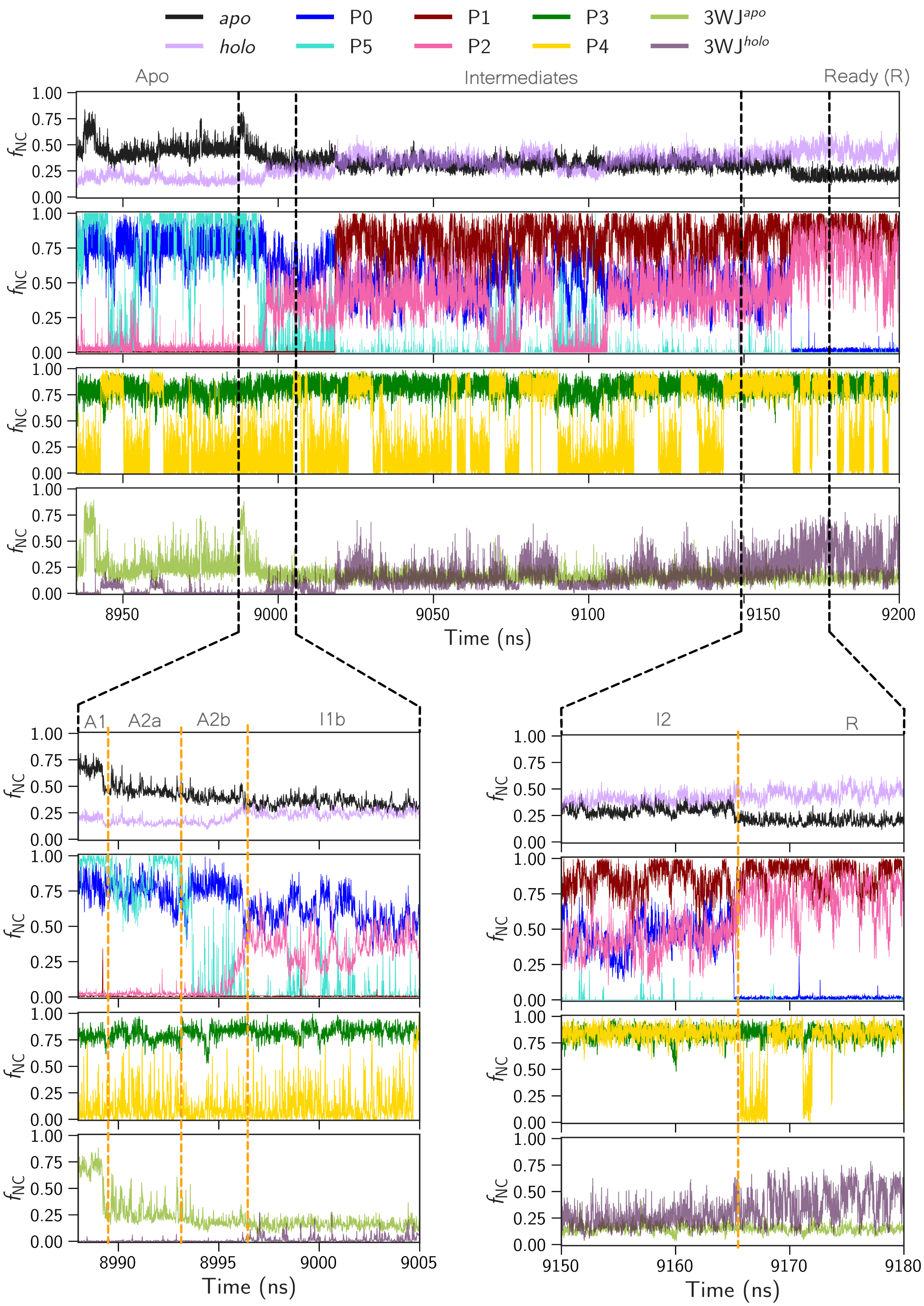


Figure S22: Representative trajectory segment for transition pathway of A1 → A2a → A2b → I1b → I2 → R. The different $f_{\mathrm{NC}}$ components are shown in four rows for transition identification.

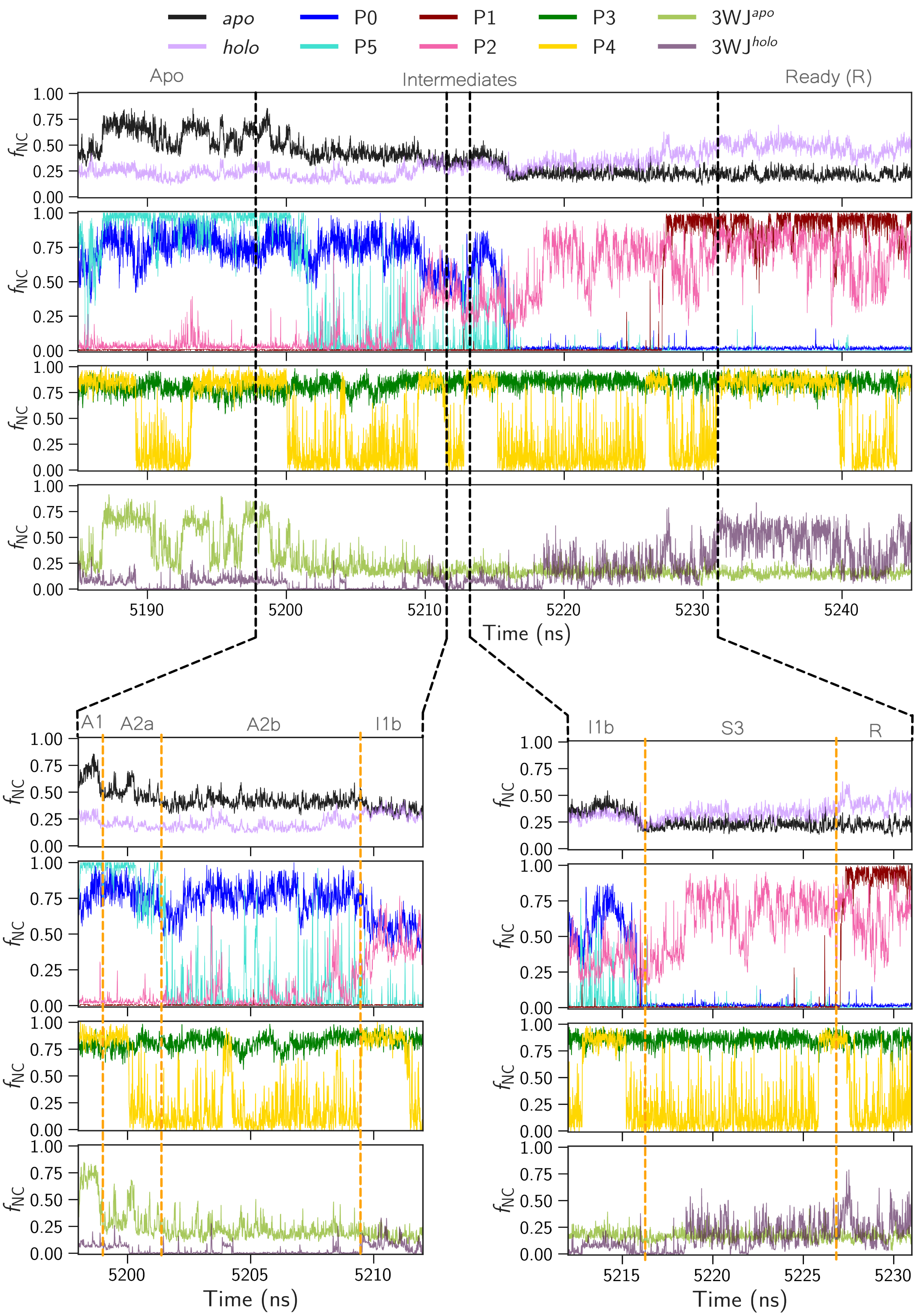


Figure S23: Representative trajectory segment for transition pathway of A1 → A2a → A2b → I1b → S3 → R. The different $f_{\mathrm{NC}}$ components are shown in four rows for transition identification.

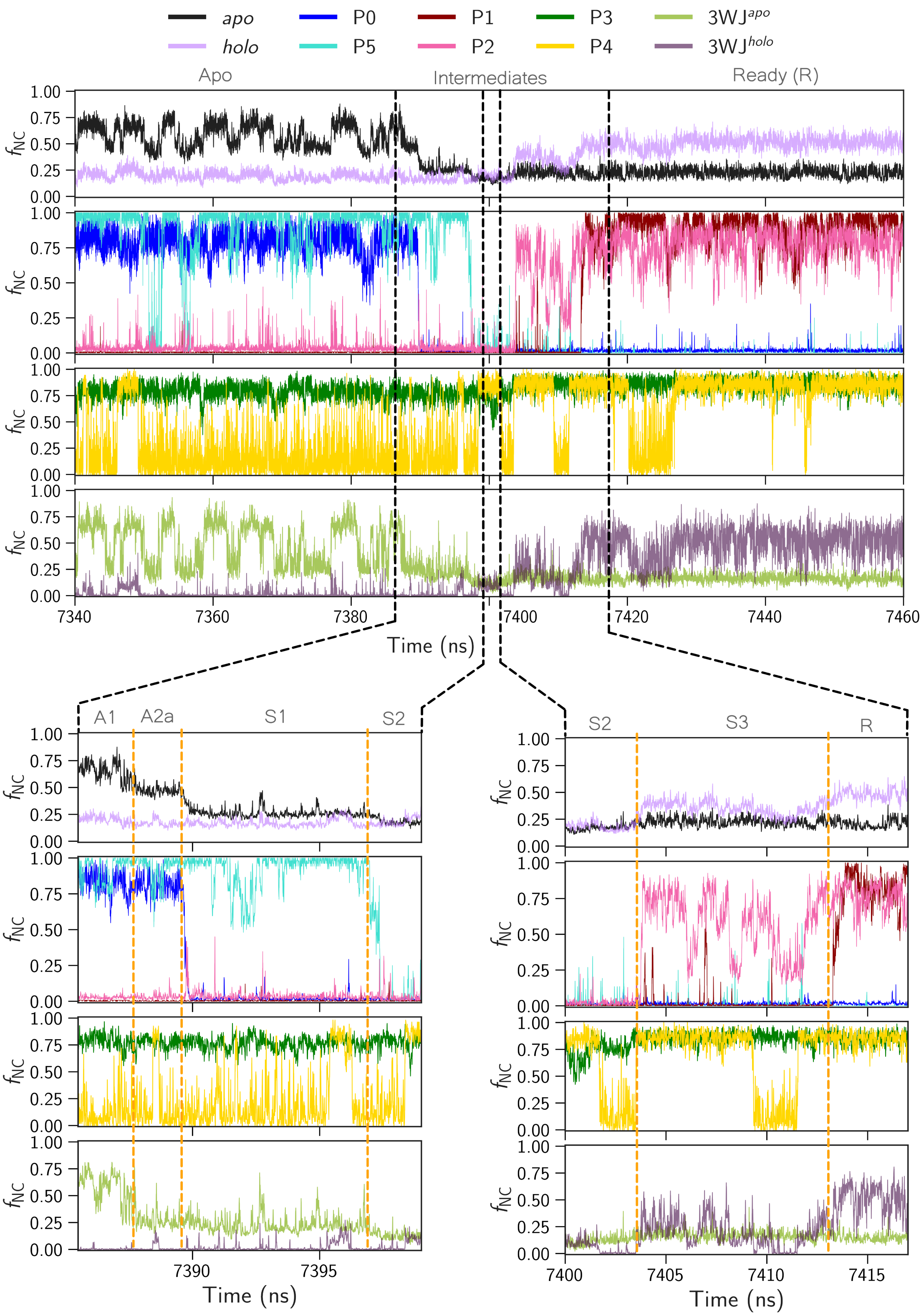


Figure S24: Representative trajectory segment for transition pathway of A1 → A2a → S1 → S2 → S3 → R. The different $f_{NC}$ components are shown in four rows for transition identification.

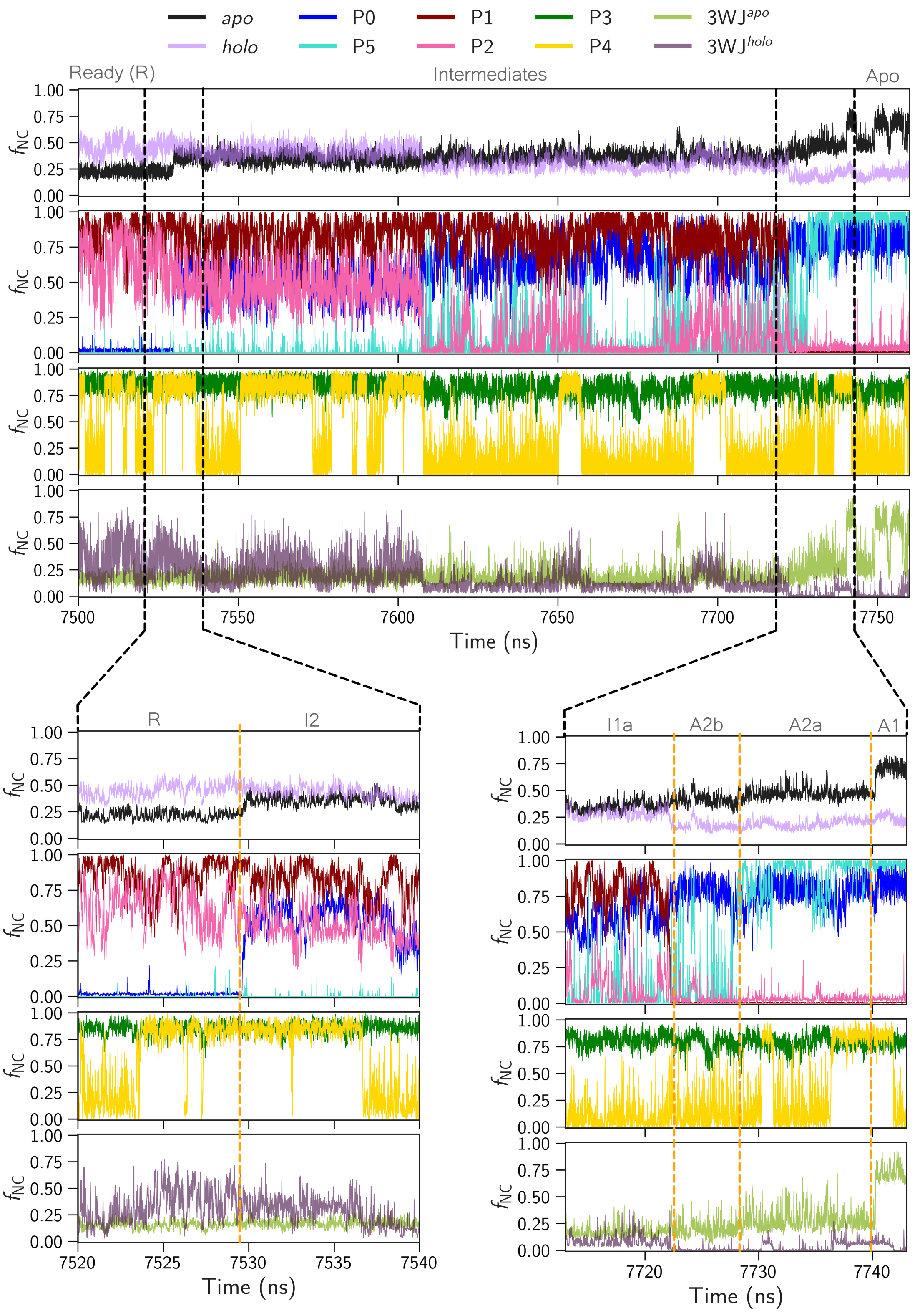


Figure S25: Representative trajectory segment for transition pathway of R → I2 → I1a → A2b → A2a → A1. The different $f_{\mathrm{NC}}$ components are shown in four rows for transition identification.

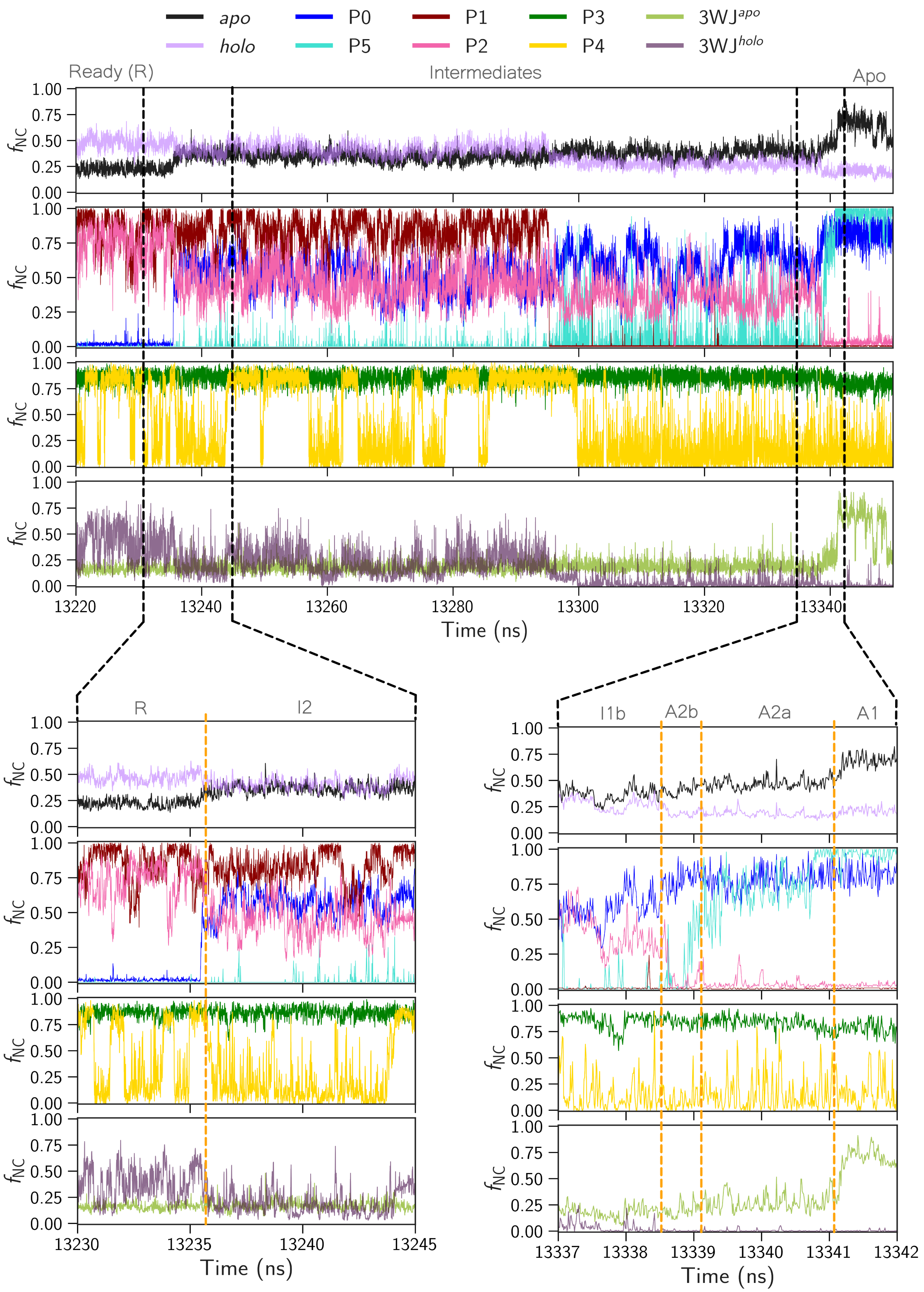


Figure S26: Representative trajectory segment for transition pathway of R → I2 → I1b → A2b → A2a → A1. The different $f_{NC}$ components are shown in four rows for transition identification.

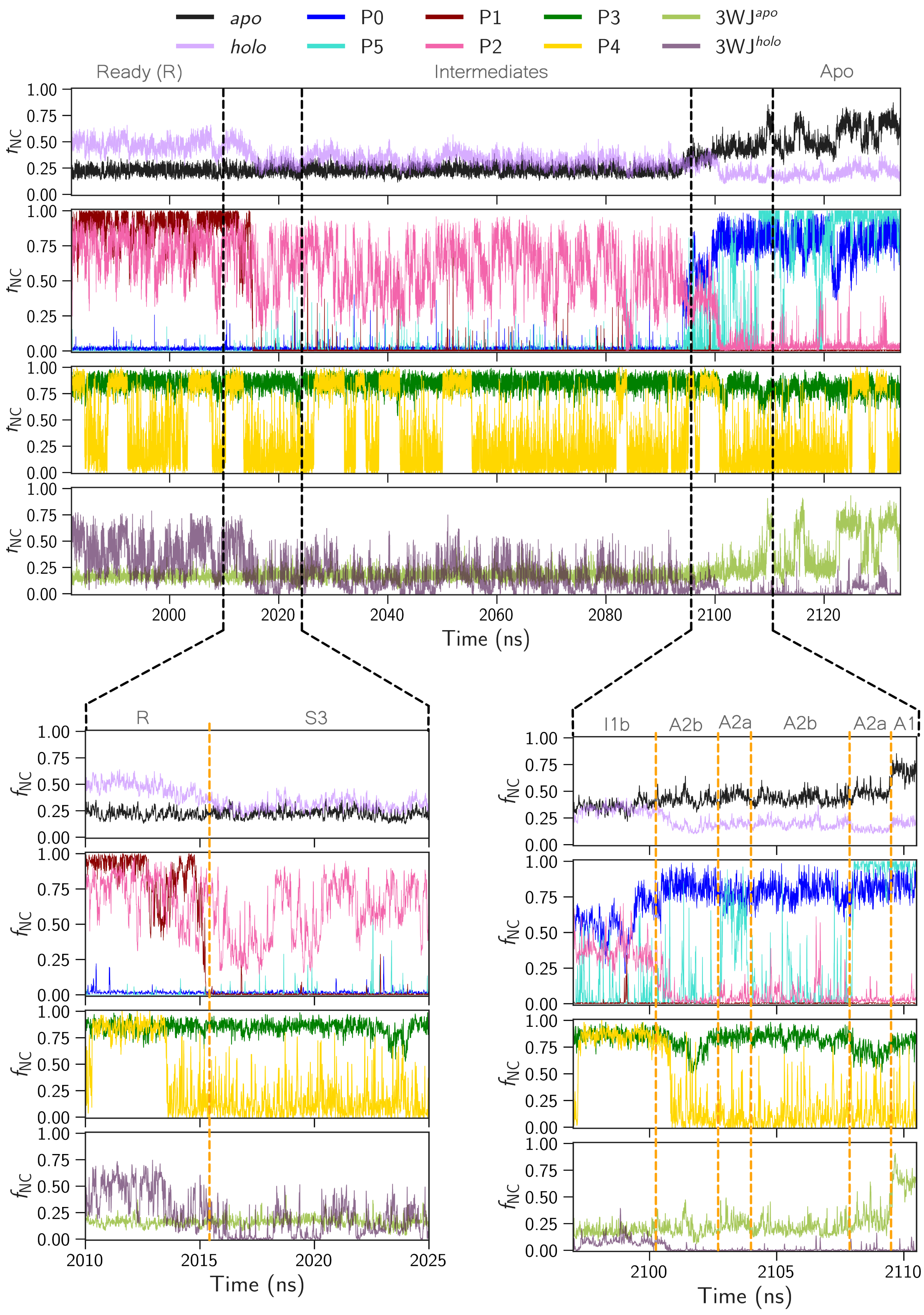


Figure S27: Representative trajectory segment for transition pathway of R → S3 → I1b → A2b → A2a → A1. The different $f_{\mathrm{NC}}$ components are shown in four rows for transition identification.

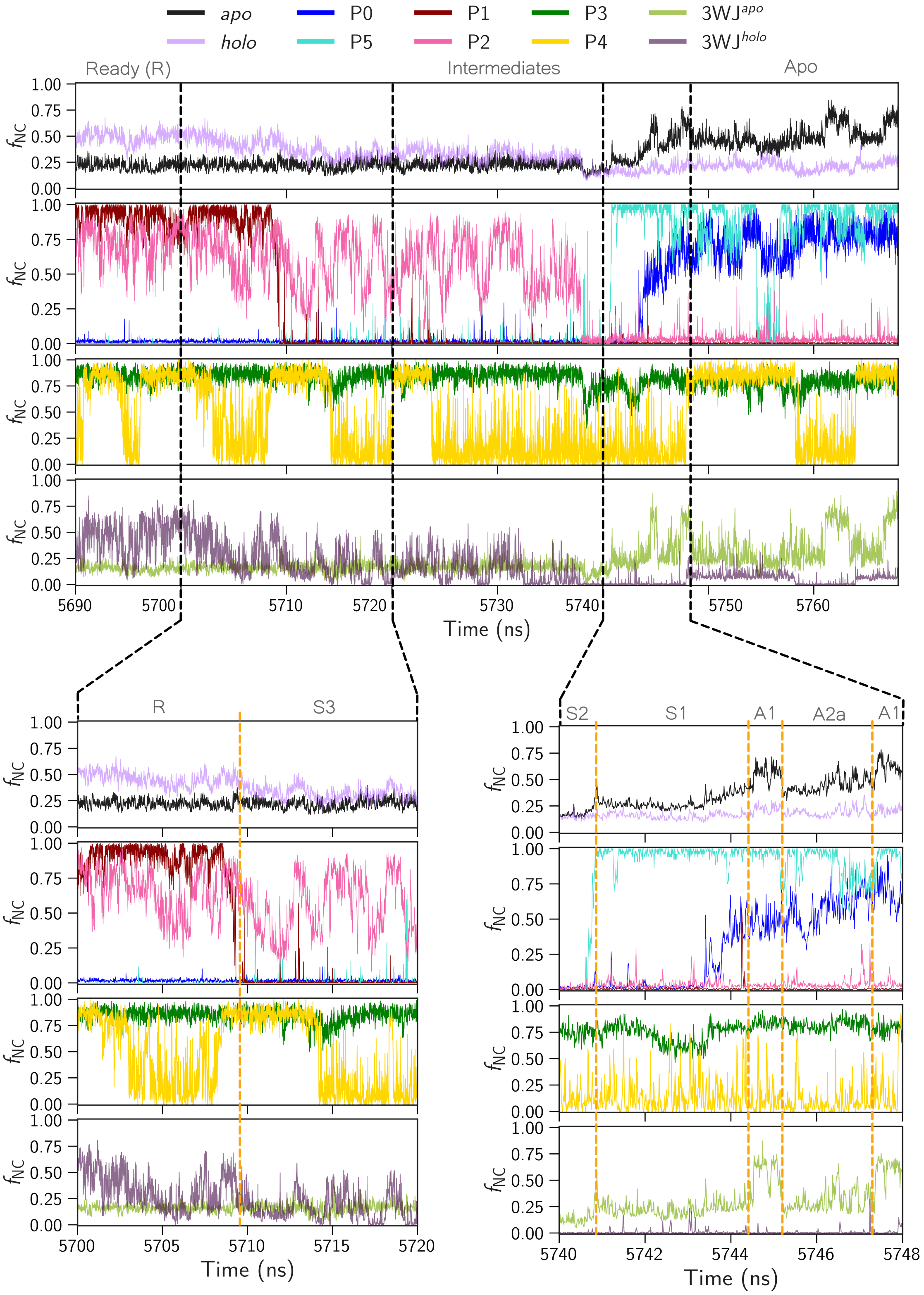


Figure S28: Representative trajectory segment for transition pathway of R → S3 → S2 → S1 → A2a → A1. The different $f_{\mathrm{NC}}$ components are shown in four rows for transition identification.
